# “Remote and partial clocks expand the circadian neuronal network, driving widespread molecular rhythmicity in *Drosophila*”

**DOI:** 10.64898/2025.12.23.696232

**Authors:** Ines L. Patop, Ane Martin Anduaga, Teddy Rashkover, Jazmin Morales, Nathan Browstein, Catherine R. Carmona, Ivana L. Bussi, M. Fernanda Ceriani, Yerbol Z. Kurmangaliyev, Sebastian Kadener

**Affiliations:** Biology Department, Brandeis University, Waltham, MA, 02454, USA; Laboratorio de Genética del Comportamiento, Fundación Instituto Leloir – Instituto de Investigaciones Bioquímicas de Buenos Aires (IIBBA-CONICET), Buenos Aires, Argentina

**Author notes:** These authors contributed equally to this work (Ines L. Patop, Ane Martin Anduaga).

## Abstract

Circadian clocks orchestrate daily physiology and behavioral rhythms, yet the extent to which cells lacking canonical clock components exhibit robust temporal regulation remains unclear. Here we addressed this question using a two-step strategy. First, we systematically mapped core clock gene expression at single-cell resolution across the *Drosophila* brain and body. Second, leveraging these data and performing targeted experiments, we uncovered cells that lack most or all canonical clock components yet display strong mRNA rhythms. We found that lamina wide-field neurons show high-amplitude *tim* mRNA cycling even in constant darkness conditions (DD) despite minimal expression of other clock genes, suggesting a partial or noncanonical oscillator. In addition, C2 and C3 optic lobe neurons, which do not express core clock components, exhibit hundreds of circadian and daily cycling transcripts. Notably, circadian rhythms in C3 neurons coincide with oscillations of activity-regulated genes (ARGs), whereas C2 neurons cycle independently of ARGs, indicating distinct non-cell-autonomous mechanisms. These findings reveal a spectrum of circadian regulation, from autonomous to remote, input-driven rhythms and expand the circadian landscape to include strategies for generating and distributing temporal information across neuronal and non-neuronal cell types.

## INTRODUCTION

Organisms have evolved circadian clocks to maintain temporal organization and predict daily environmental changes ^1^. These internal timekeeping systems are largely self-sustaining biochemical oscillators that drive rhythms in gene expression, physiology, and behavior ^2–4^. In flies, the core clock is composed of a transcriptional-translational feedback loop in specialized “clock cells” ^5^. The key players of this loop include the transcription factors CLOCK (CLK) and CYCLE (CYC), which activate the expression of other core clock genes, including *period* (*per*) and *timeless* (*tim*). The PER and TIM proteins then accumulate and eventually repress CLK-CYC activity, generating oscillations in gene expression that approximate a 24-hour period ^5–8^. This fundamental loop is further regulated by post-transcriptional and post-translational modifications, as well as additional factors like *vrille* (*vri*) and *clockwork orange* (*cwo*), ensuring accurate and robust timekeeping ^6–17^.

These molecular oscillations drive extensive downstream effects. Hundreds of mRNAs cycle in abundance across the day, coordinating numerous physiological processes and behaviors ^18–22^. The most extensively studied circadian output is the rhythm of adult locomotor activity. In *Drosophila*, as in mammals, relatively small sets of brain neurons act as primary pacemakers that generate and maintain circadian rhythms in locomotion ^5,23–25^. About 240 such clock neurons coordinate behaviors such as locomotor activity, sleep, and feeding through both direct synaptic connections and neuropeptidergic signaling. This neuronal network includes: small ventrolateral neurons (sLNv), large ventrolateral neurons (lLNv), lateral dorsal neurons (LNd), dorsal neurons (DN1–3), and lateral posterior neurons (LPN) ^24,25^. Among these circadian neurons, the ones expressing the neuropeptide *Pigment Dispersing Factor* (PDF), the sLNv and lLNv, play a central role in organizing the network ^26^. PDF in flies is functionally analogous to VIP in the mammalian suprachiasmatic nucleus (SCN, the main circadian center in the mammalian brain) ^27–29^. PDF release helps maintain robust free-running rhythms in constant darkness (DD) and shapes the prominent morning activity peak under light-dark (LD) cycles ^30,31^. Indeed, PDF is released primarily during the late night to early morning, just before lights-on ^32^. This timing and PDF’s capacity to delay evening clock neurons highlight how crucial neuropeptide signaling is for organizing behavioral rhythms under LD cycles. The PDF receptor (PDFR) is present in multiple brain regions as well as peripheral tissues^33,34^, indicating that PDF can act in both local and long-range (paracrine) manners to convey timing information beyond canonical clock cells ^35^. PDF binding to its receptor leads to increased cAMP/PKA activity which in turn stabilizes PER and TIM, slowing their degradation ^36^. This biochemical effect of PDF lengthens the period of the molecular clock and help maintain robust behavioral rhythms ^36^. Other neuropeptides, such as neuropeptide F (NPF), short neuropeptide F (sNPF), and ion transport peptide (ITP), also contribute to the circadian system in *Drosophila* ^37,38^. In recent years, advances in connectomics analyses have further deepened our understanding of this neuronal network. For instance, Reinhard et al. ^39^ employed the FlyWire connectome to identify approximately 242 clock neurons, more than the traditional 150 estimate, and discovered extensive contralateral connectivity as well as novel indirect light input pathways. These observations suggest that the circadian network is more distributed and interconnected than previously appreciated, thereby providing a robust structural framework through which central pacemakers might entrain peripheral or “clockless” cells.

Building on this distributed network architecture, neurons could convey timing information through direct synaptic contact or by releasing neuropeptides such as PDF. Indeed, calcium imaging studies of the fly brain have identified neuronal groups downstream of circadian neurons that display circadian excitation patterns peaking at different times of the day ^40^. These oscillations can be driven by individual or combined actions of various circadian neuronal groups ^41^. Among the cells exhibiting these excitation rhythms are neurosecretory cells, which release key regulatory neuropeptides such as DH44, LK, SIFa, DMS, and ILPs at specific phases of the daily cycle ^41–43^. Consistent with the idea that circadian influences extend beyond canonical clock neurons, a recent report described RNA rhythms in sorted Mushroom Body neurons from *Drosophila* brains maintained in DD^44^. However, because this work relied on bulk sorting, it is impossible to determine whether the apparent cycling arises from the MB neurons themselves or from contaminating clock-containing cells (e.g., glia). Indeed, some core clock transcripts (*per* and *Clk*) were detectable in those samples. Thus, despite these intriguing observations, it remains unresolved whether MB neurons possess autonomous molecular rhythms or instead inherit temporal signals from bona fide clock neurons.

Recent discoveries further expanded our understanding of circadian signal propagation beyond the traditional clock neuron clusters. For example, in the visual system overlapping populations of newly defined AMA (PDF-output modulatory) and xCEO (extra-clock electrically oscillating) neurons are critical for ensuring proper photoentrainment and even sustain circadian rhythms in constant conditions ^45,46^. These neurons connect to LNvs, at least one LNd and dorsal neurons projecting towards the accessory medulla, a central circadian hub, where they display ultradian electrical oscillations that modulate PDF release and help maintain coherent molecular oscillations. While their own clock status remains unclear, their extensive connections suggest that circadian timing can propagate through pathways beyond cells with canonical molecular clocks.

Within the *Drosophila* brain, glial cells harbor core clock gene oscillations and rhythmic output genes ^47–50^. In contrast, most brain neurons are thought not to contain canonical circadian feedback loops. Still, the entire brain transcriptome exhibits hundreds of cycling genes ^51,52^, suggesting a complex, multilayered control of rhythmic physiology beyond the ∼240 canonical clock neurons.

Many of these oscillations likely direct a variety of behavioral and physiological processes beyond locomotion. For example, in the lamina of the visual system, certain interneurons (L1, L2) and the glial cells surrounding the lamina cartridges display circadian changes in size and physiology ^53–56^. These structural and functional rhythms hint at underlying timekeeping mechanisms, but it is unclear whether they arise from local, cell autonomous clocks or signals from elsewhere. In particular, the presence of the PDF Receptor (*PdfR*) in some of these cells (e.g., lamina neurons ^57^), suggests that central clock neurons might impose rhythmicity on otherwise clockless cells ^58^. Remote circadian signals, including PDF and potentially other neuromodulators, could orchestrate transcriptional oscillations and physiological rhythms in cells lacking a full molecular clock. Such cells might express only a subset of core clock genes (e.g., *tim* without *per* or *Clk*) or no canonical genes at all, relying instead on extrinsic cues.

Beyond the brain, circadian rhythms extend into multiple peripheral organs in *Drosophila*, including the gut, Malpighian tubules, fat body, and reproductive organs ^59–63^. In these peripheral systems, processes such as eclosion^64^, cuticle deposition ^65^, pheromone release ^60^, mating ^66^, and odorant-driven behaviors ^67^ display rhythms. Many of these oscillations are thought to be cell-autonomous, driven by local molecular clocks. Consistent with this view, molecular and reporter-based assays have found canonical clock-gene oscillations across a broad array of peripheral sites ^53,54,56,62,68–70^. Moreover, there are hundreds of cycling mRNAs in the eye, gut, Malpighian tubules, and fat body ^58,63,70–72^. Nevertheless, we still lack precise knowledge of which specific cell types within these tissues drive these rhythmic transcripts, and whether all those cells contain a *bona fide* circadian oscillator.

Here, we employ single-cell RNA sequencing and bulk RNA-seq temporal profiling in *Drosophila*, along with systematic reanalysis of published datasets, to map core clock gene expression across tissues and identify which cell types contain canonical oscillators. We found cells that do not express all core clock components yet still show robust daily and even circadian mRNA oscillations. For example, certain lamina interneurons (Lamina wide-field-1, Lawf1 neurons) exhibit high levels of *tim* mRNA cycling under both LD and DD conditions, despite lacking other key clock genes. This finding suggests Lawf1 cells may rely on partial molecular frameworks or input from circadian pacemaker cells via neuromodulators like PDF. In addition, we found that C2 and C3 neurons in the optic lobe display robust daily and circadian gene expression without expressing clock genes. Instead, these neurons appear to depend on signals from circadian pacemakers or other upstream sources to establish and maintain their rhythmicity. Interestingly, rhythms in C2 and C3 cells arise through fundamentally different mechanisms: C3 neurons exhibit coherent oscillations in activity-regulated genes (ARGs), which continue to cycle in constant darkness, suggesting a key role for circadian modulation of neuronal activity. In contrast, ARGs in C2 neurons lose rhythmicity in DD, despite ongoing oscillations in many transcripts, indicating that these cells are driven by a distinct, ARG-independent mechanism. Together, we identify at least four distinct mechanisms for generating timing information in the *Drosophila* cells, ranging from fully autonomous clocks to entirely extrinsically-driven oscillators, thereby expanding the paradigm of circadian regulation.

## RESULTS

### Only a subset of cells across the *Drosophila* body contains a core circadian clock

To identify cell types harboring a circadian clock in *Drosophila* we used publicly available single-cell data from the brain, the optic lobe, and the whole fly ^57,73,74^. First, we mapped the expression of the main core-clock genes: *tim, per, Clk, cyc, cry, cwo,* and *vri,* focusing on *tim*, *per*, and *Clk* given that *vri* and *cwo* have clock-independent functions ^8,75–77^ and *cyc* mRNA is broadly expressed ^78^. We did so by standardizing the gene expression value of these genes across cell types and genes (Supplementary Fig. 1A, Table S1). This approach accounts for both gene-to-gene and cluster-to-cluster variability, mitigating issues stemming from sparse single-cell data and single-timepoint sampling. We validated this strategy using a brain dataset ^73^, which faithfully recovered known circadian clock-containing cells: clock neurons, photoreceptors, and glial cells (Fig.1A, top left). Applying the same method to a head dataset ^79^ we also identified cell types known to have a circadian clock: eye and photoreceptor cells, multiple glial types, and fat body cells; circadian neurons were not recovered, likely due to insufficient cell coverage of the head dataset ^79^ (Supplementary Fig. 1B).

**Figure 1.**
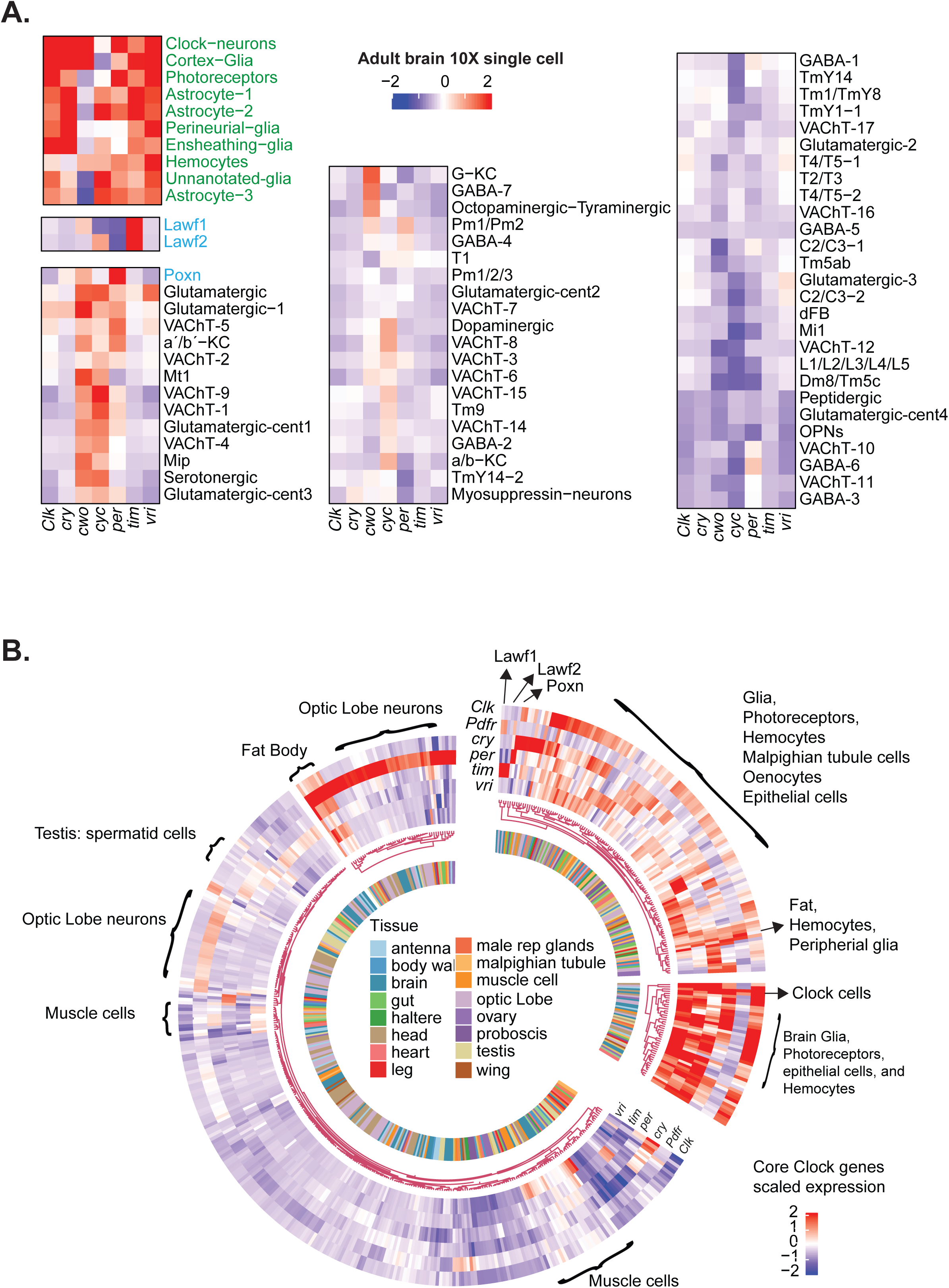
Only a subset of cell types express core clock components in *Drosophila.* **A.** Heatmap of standardized core-clock gene expression across cell types from 10X single-cell data *Drosophila* brain. The red and blue color scale represents the relative centered levels (>0 or <0 respectively) (data from ^73^). **B.** Heatmap of standardized expression values across cell types in the whole *Drosophila body* clustered by similarity (integrated data from ^57,73,74^). Blue and red represent decreased and increased expression, respectively. Colors in the inner part represent the tissue of origin for each cluster. The main cell types of interest are identified individually.

Next, we extended our analysis to the rest of the fly body using the fly cell atlas and other single-cell datasets ^57,73,74^ (Supplementary Fig. 2, Supplementary Fig. 3, Table S1). We observed core clock gene expression in various cell types, including oenocytes, hemocytes, and epithelial cells, across tissues like the body wall, gut, Malpighian tubules, leg, heart, wing and halters (Supplementary Fig. 2A-H). Notably, these peripheral datasets revealed no clock gene expression in neurons, yet epithelial, tracheal, and glial cells commonly expressed clock genes. Interestingly, muscle tissues displayed differential core gene expression: while antenna muscles expressed most core clock genes, other muscle groups lacked *Clk*, *tim*, or *per*, strongly suggesting the absence of an endogenous circadian clock (compare Supplementary Fig. 3A with panels in Supplementary Figs. 2 and 3). Surprisingly, we detected no core-clock gene enrichment in olfactory receptor neurons of the antenna or gustatory receptor neurons of the proboscis (Supplementary Fig. 3A, B), despite reports of cycling physiology ^56^. In male reproductive tissues, we found clock gene enrichment in cells of the seminal vesicle, ejaculatory bulb, and duct (Supplementary Fig. 3C, D) as previously reported ^80^. In ovaries (Supplementary Fig. 3E), we observed *tim* and *per* expression in follicle cells, but not in oocytes or nurse cells, consistent with previous findings ^81^.

We then integrated the data from all tissues, scaling values, incorporating the receptor for the neuropeptide PDF (*Pdfr*) and clustering the cells based on the expression of the clock genes (Fig. 1B). Importantly, cell types found in multiple datasets (e.g., optic lobe neurons which are present in brain, optic lobe, and head datasets) clustered together, validating our integration. This global integration distinguished cells enriched or depleted for core clock genes (Fig. 1B). Approximately one-third of cell types expressed high levels of most clock genes (*Clk*, *per*, *tim*, *vri*, or *cry*). This subset included glia, circadian neurons, epithelial cells, all Malpighian tubule cells, oenocytes, fat cells from various tissues, and hemocytes. Within this “clock-enriched” group, some cells expressed only one or a few clock genes (Fig. 1A, B and fig.S2B). For example, we found that Lawf1 neurons (lamina wide-field 1 interneurons in the visual system) express high levels of *tim* but not *per* or *Clk*, suggesting a partial clock. Conversely, Poxn neurons (a subset of neurons that express *Poxn*) express *per* without *tim* or *Clk*. In addition, most muscle cells express only *vri.* While *vri* is known to have circadian-independent functions in the heart muscle ^82^ and hair cells ^83^, the unexpected high levels of *tim* in Lawf neurons and *per* in Poxn neurons were striking. Overall, ∼65% of cell types lacked enrichment for any core clock genes (Fig.1B, fig S2, S3 and Table S1), underscoring that most neurons, especially in the brain, lack robust core clock gene expression. Nevertheless, many clockless cells express *PdfR*, suggesting these cells may still receive circadian signals through neuromodulators like PDF (Fig.1B and see below).

### *Tim* is highly expressed and display circadian changes in Lawf neurons

To quantify the contribution of each cell type to overall clock gene expression in the brain, we calculated the proportion of total core-clock gene signal per cell type (Table S2). As expected, glia served as the main contributor, and particularly the ensheathing glia-comprising 6% of brain cells-accounted for 23% of total *tim* expression. Interestingly, smaller clusters, such as clock neurons, Poxn, and Lamina wide-filed 1/ 2 (Lawf1/2) neurons, also contributed significantly to *Clk*, *per*, and *tim* signals, respectively (Table S2, Figures 1B, 2A and Fig.S4A). In particular, Lawf1 neurons showed the highest *tim* enrichment in the fly brain, surpassing all glial subtypes and known clock neurons (Fig. 2A, Table S1, S2). However, we did not detect other clock genes like *Clk* or *per* in these cells (Fig. 2A, Supplementary Fig. 4A and Table S1, S2).

**Figure 2.**
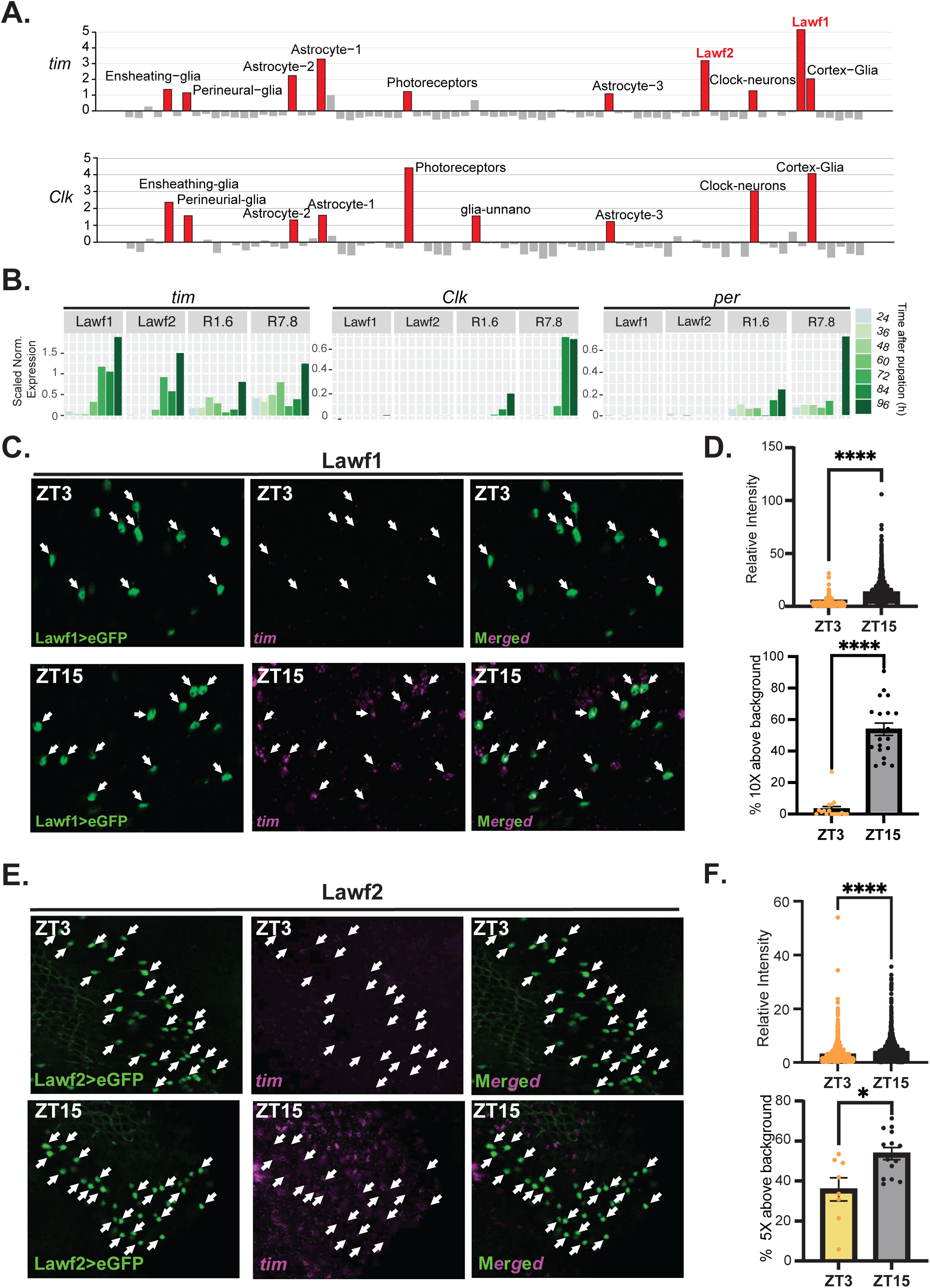
*tim* is highly expressed and display daily changes in Lawf neurons. **A.** Standardized *tim* (top) and *Clk* (bottom) expression from 10X single-cell data from the brain. In red, cell types with expression levels higher than one standard deviation from the mean (from ^73^). **B.** Bar plot showing the log normalized expression of *tim, Clk* and *per* in the indicated cell types during pupal development (from ^57^). **C.** Representative images of *tim* mRNA expression in Lawf1 cells at ZT3 (top) and ZT15 (bottom). Cells were visualized by GFP i(Lawf1-Gal4; UAS-eGFP flies, left panel). *tim* expression by *in situ* hybridization using probes against *tim* coding region (center). Right overlay of the two images. **D.** Quantification of the experiment in C. Top: quantification of *tim* mRNA fluorescence intensity in individual Lawf1 neurons at the indicated timepoints. Bottom: proportion of Lawf 1 neurons per optic lobe with *tim* signal ≥10-fold above background. Mann Whitney test, **** p <0.0001. **E.** Representative images of *tim* mRNA expression in Lawf2 cells ZT3 (top) and ZT15 (bottom). Cells were visualized by GFP (Lawf2-Gal4; UAS-eGFP flies, left). Center: *tim* expression by *in situ* hybridization using probes against *tim* coding region. Right: overlay of the two images. **F.** Quantification of the experiment described in E. Top: quantification of *tim* mRNA fluorescence intensity in individual Lawf2 neurons. Lawf2 neurons identified by GFP expression driven by Lawf2-GAL4. Bottom: proportion of Lawf 2 neurons per optic lobe with *tim* signal ≥5-fold above background. Mann Whitney test, * p<0.05, ****p <0.0001.

We confirmed these findings using another single-cell dataset ^74^ and additional optic lobe bulk RNA sequencing data from single cells and sorted populations ^57,73^ (Supplementary Fig. 4B and Table S1, S3). *Per*, *Clk* and *vri* levels are extremely low in Lawf cells (Table S1-S3, Fig. 2A and fig.S4A) suggesting these genes are not expressed in these neurons. Moreover, *tim* but not *per* or *Clk*, shows strikingly high expression during pupal development in Lawf1 and Lawf2 neurons, exceeding the levels found in other cells that harbor a circadian clock like R1-6 and R7-8 photoreceptors (Fig. 2B) ^57^. This result suggests that *tim* expression relies on a different, likely CLK-independent, mechanism for expression in these neurons. In this context, the presence of high *tim* levels but no other key circadian components point to a non-canonical timing mechanism for *tim* expression in Lawf cells.

To test whether *tim* levels exhibit daily oscillations in Lawf cells, we measured *tim* mRNA levels in Lawf1 and Lawf2 neurons at ZT3 and ZT15 (trough and peak *tim* expression, respectively) in flies maintained under 12:12 Light: Dark (LD) conditions using *in situ* hybridization. We identified Lawf1 or Lawf2 neurons by using well-characterized drivers (see methods) to drive expression of a green fluorescent protein (GFP) reporter. As expected, the GFP signal shows that Lawf1 and Lawf2 cell bodies reside in the medulla cortex and project towards the medulla and lamina neuropils (76–78) (Supplementary Fig. 4C). Importantly, *in situ* hybridization revealed robust, high amplitude *tim* oscillations in Lawf1 neurons (Fig. 2C, D and Supplementary Fig. 4D). Lawf2 also expressed *tim*, but at lower levels than Lawf1. Interestingly, not all Lawf2 cells show *tim* signal, at least in the assayed timepoints (Fig. 2E, F and Supplementary Fig. 4E), which aligns with the single-cell data in which only a fraction of Lawf2 neurons express *tim* (Fig. 2A, Table S2). We also observed significant changes in *tim* expression in Lawf2 cells between the two timepoints (ZT3 and ZT15), but these changes were smaller than those in Lawf1 cells (Fig. 2E-F, Supplementary Fig. 4E, Table S3). To explore this further, we assessed *tim* expression in Lawf2 cells using a *tim*-Tomato transcriptional reporter from our lab that peaks in the late night/early morning (12). We found that the *tim-*Tomato reporter is expressed in Lawf2 cells (Supplementary Fig. 4F) and that the levels fluctuate over the course of the day in these cells (Supplementary Fig. 4G, H), which corroborates the *in-situ* results and demonstrates that Lawf2 neurons harbor an oscillator that drives rhythmic *tim* transcription.

### Lawf1 neurons express *tim* in a circadian fashion and are connected to cells with a circadian pacemaker

To determine whether these oscillations are truly circadian, we assessed *tim* mRNA oscillations in Lawf cells under constant darkness (DD). We focused on Lawf1 neurons because they show a stronger *tim* signal in single-cell sequencing and *in situ* hybridization, as well as high-amplitude oscillations under 12:12 Light: Dark (LD) conditions (see above). In agreement with the LD data, strong changes in *tim* levels emerged between the two assayed timepoints in Lawf1 neurons during the first day in DD (DD1), with high *tim* levels at CT15 (Circadian Time 15) and lower at CT3 (Circadian Time 3, Fig. 3A, B and Supplementary Fig. 4I). These results show that *tim* mRNA cycling persists in constant conditions and confirm that Lawf1 neurons exhibit true circadian regulation despite lacking a canonical oscillator.

Because Lawf1 cells do not harbor a full molecular clock, we hypothesized that they must receive timing information from other cells. To identify plausible sources, we examined the recently published *Drosophila* visual system connectome ^84^. This analysis revealed synaptic pathways linking Lawf1 cells to both circadian neurons (l-LNvs) and photoreceptors (Fig. 3C), two cell types with canonical clocks and strong rhythmic gene expression. More specifically, Lawf1 cells receive indirect input from photoreceptors and l-LNvs via L3 and aMe4 neurons respectively (Fig. 3C). Moreover, Lawf1 is one of the principal synaptic targets of the l-LNvs although with relative few direct contacts. AMA neurons (including aMe12 ^45^) also participate in these circuits. Lawf1 neurons (but not Lawf2) express *Pdfr* (Fig. 3D), suggesting that l-LNvs could relay timing information through PDF either directly or via paracrine signaling. Additional neuropeptides and/or neurotransmitters could provide timing cues to Lawf2 neurons as well as further modulate Lawf1 circadian gene expression (Fig. 3D).

**Figure 3.**
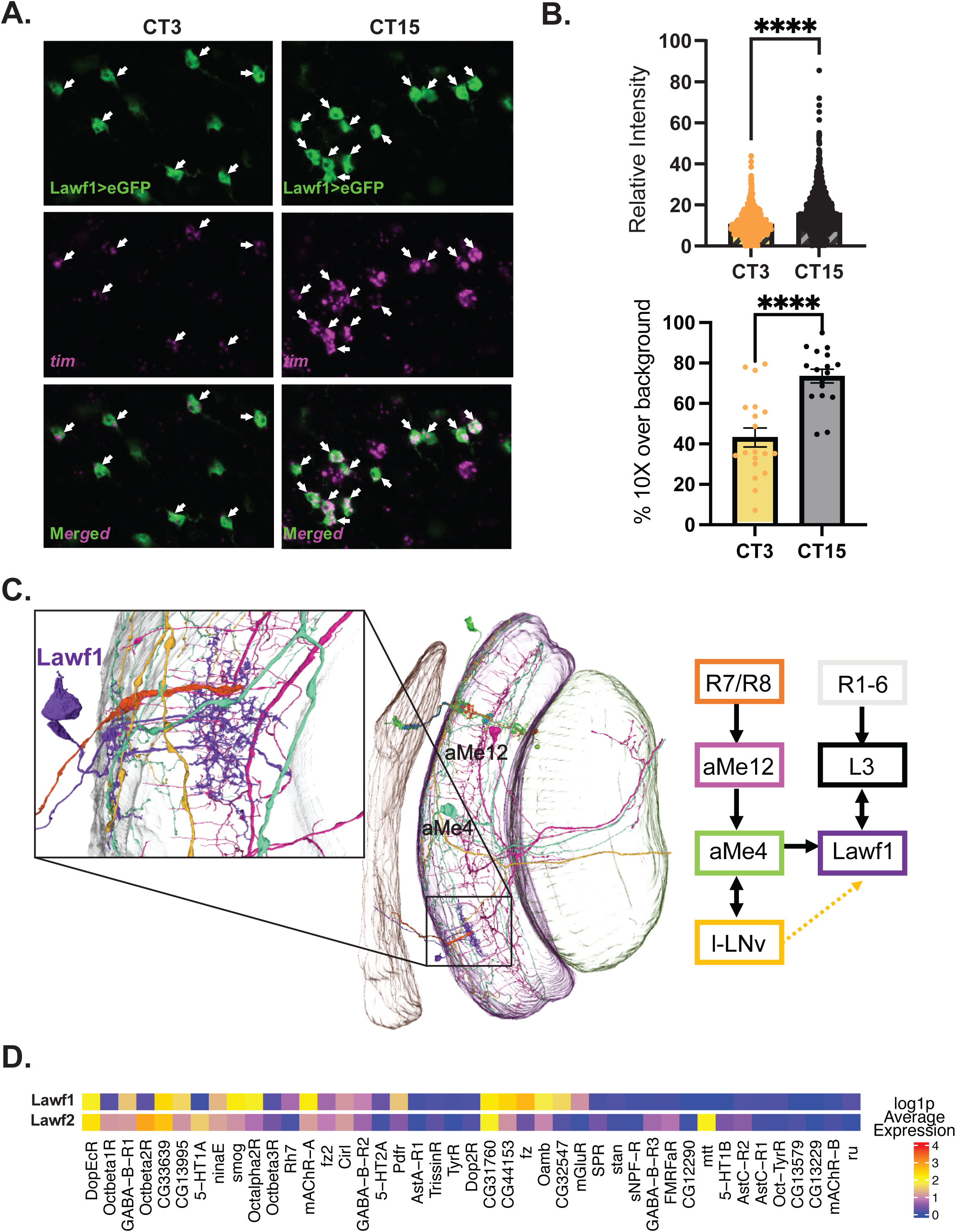
Lawf1 cells display circadian *tim* expression. **A.** Representative images of *tim* mRNA expression in Lawf1 cells at two different circadian timepoints in DD1 (CT3 and CT15). Top: Lawf1 cells. Cells were visualized by GFP in Lawf1-Gal4; UAS-eGFP flies at the indicated timepoints. Middle: *tim* expression as visualized by *in situ* hybridization using probes against *tim* coding region. Bottom: Overlay of the two images. **B.** Quantification of the experiment described in A. Top: quantification of tim mRNA fluorescence intensity in individual Lawf1 neurons at two circadian timepoints (CT3 and CT15). Lawf1 neurons identified by GFP expression driven by Lawf1-GAL4. Bottom: proportion of Lawf 1 neurons per optic lobe with tim mRNA signal ≥10-fold above background. Significance assed using Mann Whitney test, **** p <0.0001. **C.** Connectomic linkage of photic and circadian pathways to Lawf1 and Lawf2 cells. Left: visualizations of Janelia optic lobe connectome v1.1 EM-reconstruction marking the lamina (brown), medulla (purple), accessory medulla (faint purple) and lobula (green). Selected neurons capture converging photic and circadian synaptic pathways. Right: Simplified connectome of Lawf1 and lLNV neurons. Strong connections are shown between the selected cell types and Lawf1 and l-LNV neurons ^84^. **D.** Heatmap of GPCR gene expression (GPCRs from ^92^) of highly expressed transcripts in select optic cell types using the dataset ^57^.

Together, these findings show that Lawf neurons can exhibit circadian gene expression without an intact intracellular oscillator, implying that non-cell-autonomous inputs sustain their rhythms. This prompted us to ask whether other brain cells entirely lacking core clock components also display circadian changes in gene expression.

### Many mRNAs displaying daily cycling are enriched in cells without a core circadian oscillator

We therefore broadened our analysis to determine how widespread rhythmic mRNA expression is among brain cell types that lack all canonical clock genes. Previous work identified hundreds of oscillating RNAs in diverse fly tissues, including heads and brain ^51,63,85–87^. To identify robust cycling genes in the brain, we generated a new RNA-seq dataset from control *w*^1118^ brains collected every four hours in LD, identifying 231 strong cyclers. Most of these genes (219) show robust expression in single-cell sequencing data (Table S4). We then mapped them onto a single-cell dataset ^73^. Roughly half of the cycling genes are highly enriched in glia, hemocytes, and photoreceptors, cell types known to harbor canonical clocks (see clusters 1–3 in Fig. 4A and Tables S4, S5). This pattern suggests that their oscillations arise from classic cell-autonomous mechanisms.

**Figure 4:**
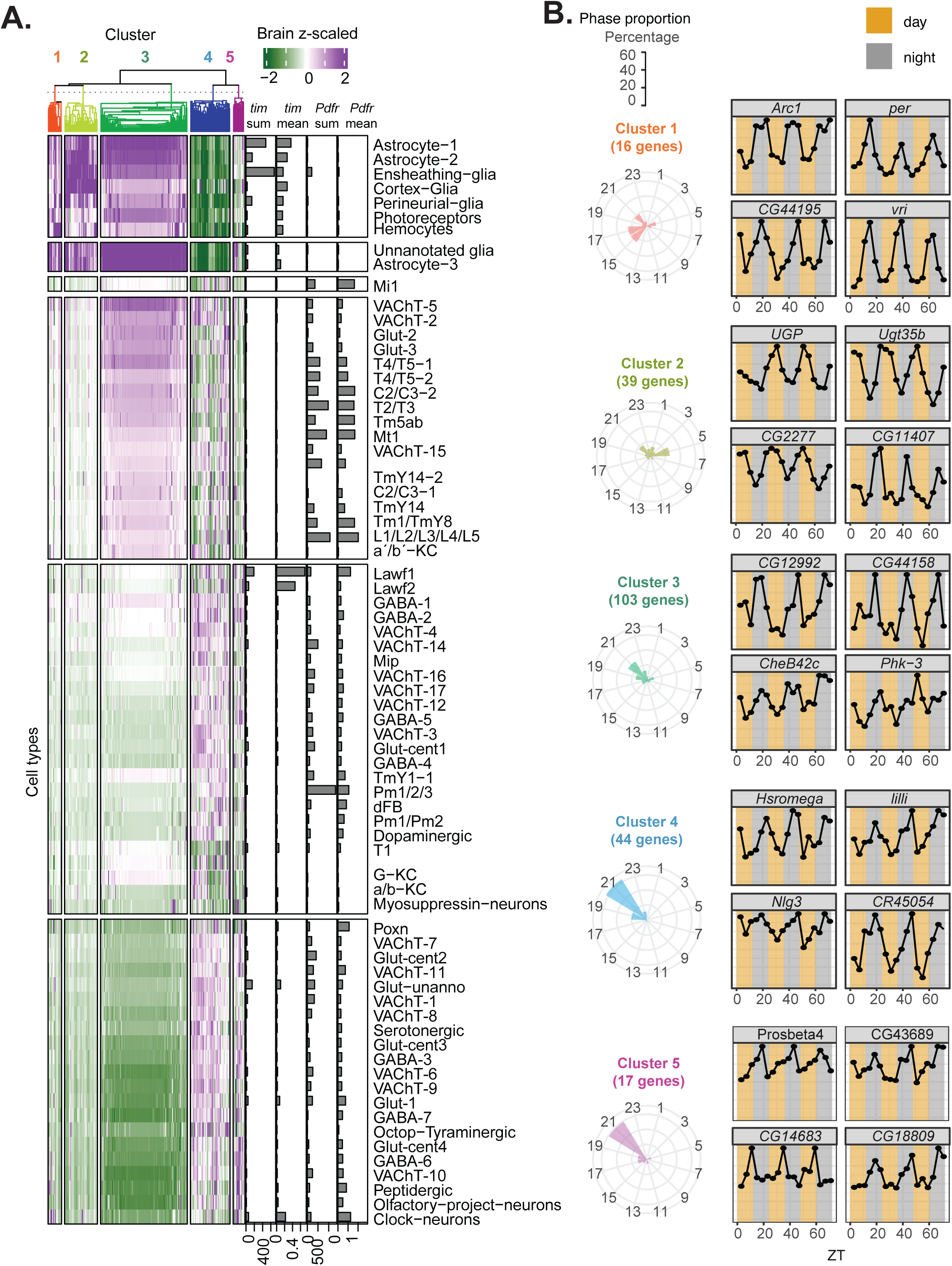
Genes with cyclic expression in the brain are enriched in cells with or without core-clock expression. **A.** Heatmap of standardized expression values by cell type clustered by similarity. Bar graph shows sum and mean expression of *tim* and *Pdfr* for each cluster. **B**. Left panel: Proportion of genes by cluster cycling at different phases. Right panel: Cycling of representative genes. Expression is normalized by the maximum (n=3, 30 flies per replica per timepoint, FDR<0.05 and amplitude > 1.5) and phase is calculated using JTK algorithm.

In contrast, the remaining genes are enriched in brain cell types with no detectable expression of core clock components (clusters 4 and 5 in Fig. 4A), implying that their rhythms may be driven by alternative mechanisms or by timing cues relayed from central pacemakers. Because our our approach reliably detected clock genes in all established clock-harboring cells, the lack of clock gene expression in these populations likely reflects true absence rather than technical dropouts.

To determine whether this pattern extends beyond the brain, we examined two peripheral tissues, gut and Malpighian tubules, for which temporal RNA-seq datasets are available ^71^(Supplementary Fig. 5A and B respectively). In Malpighian tubules, all cells express core clock components, indicating that cycling expression in this tissue is driven by canonical clocks (Supplementary Fig. 5A). However, in the gut, subsets of cycling mRNAs map to cells lacking clock genes, such as the visceral muscle cells in the crop and midgut (Supplementary Fig. 5B), indicating that “remote” regulation of daily mRNA oscillations also occurs in peripheral tissues. Together, these findings support a model in which rhythmic gene expression can arise independently of cell-autonomous clocks across multiple contexts, pointing to widespread remote circadian regulation throughout the fly.

Circadian neuropeptides may mediate these remote signals. Indeed, many clockless cells that enriched for cycling mRNAs express high levels of *Pdfr*, including several neuronal cell types in the brain and visceral muscle cells in the gut (see Fig. 4A and Supplementary Fig. 5B). Interestingly, analysis of *tim* and *Pdfr* expression revealed that most cells express either *tim* or *Pdfr*, but not both (Fig. 1B, Fig. 4A, Supplementary Fig. 5C, and Table S1), suggesting that PDF may drive oscillations specifically in cells lacking canonical molecular clocks. A small number of cell types, including certain glia, fat cells, clock neurons, and Lawf1 neurons, express both *tim* and *Pdfr* (Supplementary Fig. 5C). In these populations, PDF may adjust the oscillator phase (fat cells), promote synchrony (clock neurons), or even drive *tim* oscillations (Lawf1). Moreover, within each “cell-type cluster”, cycling genes peak at similar phases. Notably, the cycling genes enriched in high-*Pdfr* clusters (4 and 5) peak primarily around ZT 21 (Fig. 4B) coinciding with the peak of Ca^2+^ signaling in sLNVs ^88^, further suggesting that PDF may modulate these oscillations.

Altogether, these patterns indicate that extensive rhythmic transcription can occur in brain cell types that lack complete molecular clocks, prompting us to examine this phenomenon within specific neuronal populations.

### C2 and C3 neurons exhibit widespread and coordinated daily oscillations in gene expression

Building on the observation that many cycling transcripts arise in cells lacking canonical clock components, we sought a neuronal population in which we could experimentally test this phenomenon with high temporal resolution. C2 and C3 interneurons provided an ideal opportunity because they are completely clockless, as demonstrated by the absence of *Clk*, *cyc*, *tim*, *per*, and *vri* mRNAs (Fig. 1, Supplementary Fig. 1B and Data□S1-S3). Importantly, these neurons are molecularly well-defined and transcriptionally homogeneous^89^, making them an attractive system in which to assess rhythmicity at single-cell resolution. In addition, C2/C3 cells can be isolated in sufficient numbers for temporal profiling, enabling a direct test of whether coordinated daily transcription can emerge without an intrinsic molecular oscillator.

C2 and C3 neurons are also embedded within circuits connected to cells harboring a canonical circadian clock. Based on data obtained from the optic lobe connectome^84^, we found that both neuron types receive dense synaptic input from lamina monopolar cells L1–L5 (Fig.□5A). In addition, Mi1 neurons, which receives output from l□LNvs, ranks as the second largest presynaptic partner of C2 and the fourth□largest of C3 (Fig.□5A and ^84^), providing a plausible anatomical route for circadian signals. In addition, these cells express several G□protein□coupled receptor, some of which were previously described as circadian-related.□For example, C2 and C3 express RNAs encoding *AstA*□*R1* and *DopEcR*; C2 expresses the mRNA encoding the serotonin receptor *5HT*□*1A*; octopamine□receptor distribution differs, with *Oct*α*2R* in C2, *Oct*β*2R* in C3 and *Oct*β*1R* in both types (fig.□S6A). The neuropeptides and neurotransmitters activating those receptors have been previously implicated in some aspects of circadian rhythms and sleep ^90–93^.

To test for mRNA rhythmic mRNA oscillations in C2/C3 neurons, we performed single-cell RNA sequencing of these neurons at six circadian time points in light-dark (LD). and the first day in constant darkness (DD1). We used a genetic multiplexing technique ^89,94^, for pooled profiling of all time points in a single experiment to reduce technical variation between samples (Supplementary Fig. 6B). As expected, core clock mRNAs remained undetected (*Clk, tim, vri, cyc, cry*) in both neuron types confirming that these cells lack a canonical circadian molecular machinery (Figure□5B, Data□S6). *per* was absent in C2 and barely above threshold in <10□% of C3 cells on average, below levels seen in *bona-fide* clock-containing populations ^48^. *Cwo* showed low but consistent signal in C3 (LD and DD) and marginal expression in C2 (LD only). For cycling analysis in this single cell data, we split each timepoint into three pseudo replicates (as in ^48^, Supplementary Fig. 7B) and used the JTK^95^ and the harmonic mean method ^96^ followed by FDR to identify cycling genes. *per* not only is lowly expressed but does not display oscillations in LD or DD (Supplementary Fig. 6C). *Cwo* showed a robust LD oscillation in C3 that peaked at midday, opposite to its phase in canonical clock cells ^6–8,17^, and lost rhythmicity in DD (fig.□S6C). Despite the absence of a canonical oscillator, we identified 359 cycling mRNAs in C2 and 301 in C3 neurons under LD conditions (Fig.□5C; Data□S6). Their phases clustered tightly around midday (ZT6) and midnight (ZT18), indicating coordinated transcriptional programs (Fig.□5C). Cycling mRNAs included activity regulated genes (ARGs) in both cell types (see below), as well as mRNAs encoding transcription□factors (e.g. *CrebA* and *fru* in C3 neurons), numerous mitochondrial and ribosomal□protein mRNAs and other regulatory proteins (Fig.□5D; fig.□S6D and Table S6).

Given that PDF conveys morning signals ^35,97^ and that photoreceptors respond to light, we examined ARG expression in our datasets. We focused on the 4 stronger *Drosophila* ARGs (*Hr38, sr, CG14186* and *CG46385*), defined as those that have been found to be differentially expressed in brains exposed to at least 2 of 3 acute activation paradigms ^98^. All four cycled with high amplitude and similar phase in both C3 and C2 cells under LD (Fig. 5D, Supplementary Fig. 6D and Table S7). We then assessed the broader ARGs set and found that the aggregate signal of all cycling ARGs (12 in C2; 10 in C3) showed coherent mid-day peaks with pronounced oscillations and highly correlated waveforms across transcripts in both neuron types (Fig. 5E and Supplementary Fig. 6E).

**Figure 5:**
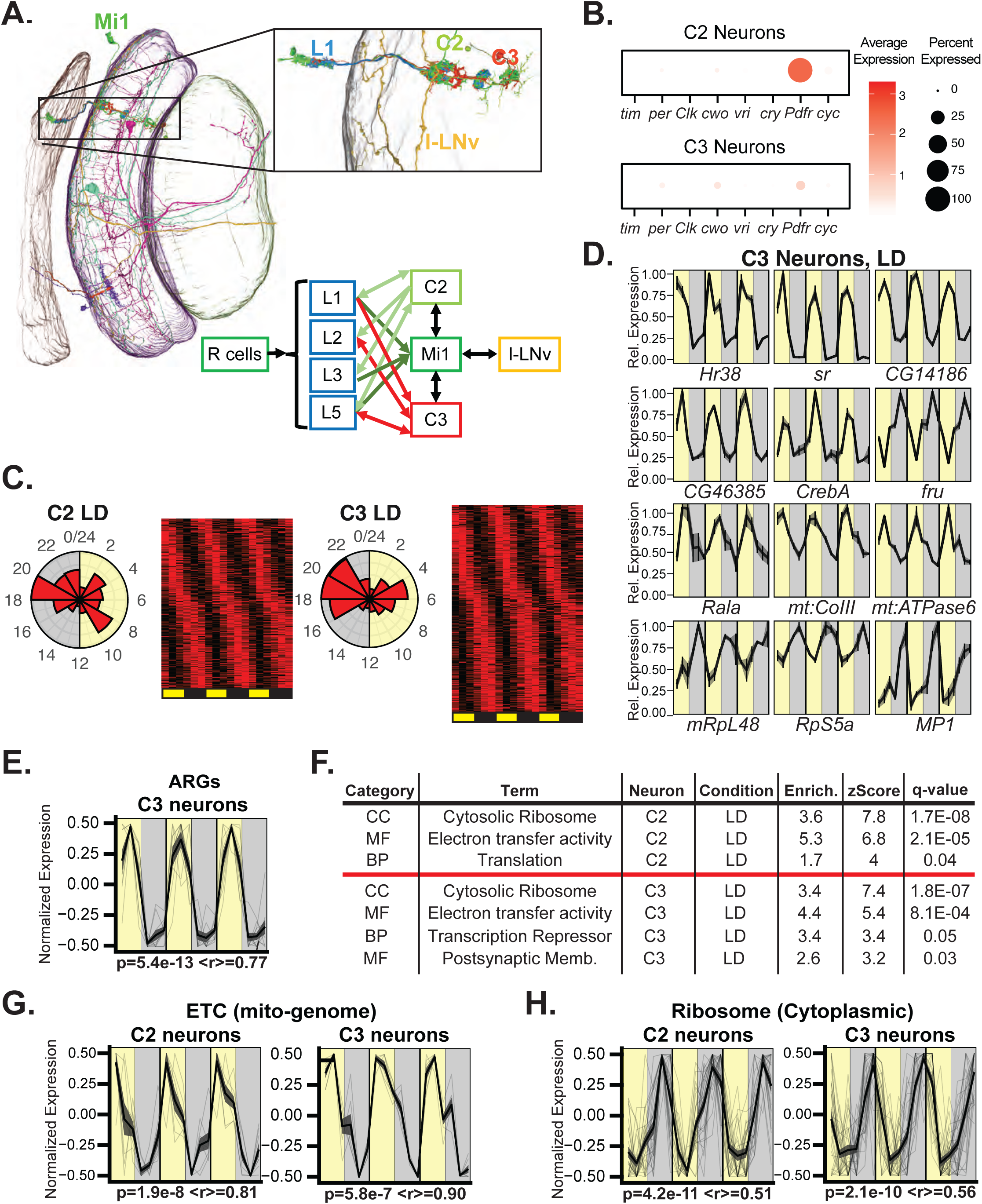
C2 and C3 neurons exhibit rhythmic and coordinated transcript expression. **A.** Connectomic linkage of photic and circadian pathways to C2 an C3 neurons. Left: Visualizations of optic lobe connectome v1.1 EM-reconstruction marking the lamina (brown), medulla (purple), accessory medulla (faint purple) and lobula (green). Right: Simplified connectome of C2/C3 and l-LNV neurons. Strong connections are shown between the selected cell types and C2, C3 an l-LNv neurons. **B.** Expression of core circadian genes and *Pdfr* in C2 and C3 neurons. Data from all timepoints for C2 or C3 neurons was averaged. **C.** Phase plot (left) and Heatmap (right) of cycling genes using scaled gene expression across times in C2 and C3 neurons in LD. **D.** Cycling gene profiles in C3 LD, plotted as max-normalized means with S.E.M. among pseudoreplicates. **E.** Aggregate waveforms of cycling ARG transcripts in C3 neurons in LD conditions. Faint lines mark individual genes, dark lines indicate signal averages, ribbons mark S.E.M. The waveforms were rescaled to -0.5 to 0.5 by subtracting 0.5. p indicates meta2d p-value and r refers to the average of pairwise Pearson’s correlations between the individual genes and the average waveform **F.** Strongest and representative Gene Ontology enrichments for the LD rhythmic mRNAs in C2 and C3 **G, H.** Aggregate waveforms of cycling ETC transcripts encoded by the mitochondrial genome (G) or cycling RNAs encoding cytoplasmic ribosomal proteins (H) in C2 or C3 neurons in LD conditions. The waveforms were generated and tested as described in E.

Gene Ontology (GO) enrichment analysis of LD-cycling transcripts revealed significant overrepresentation of cytosolic ribosomal components and electron transport chain (ETC) genes in both C2 and C3 neurons (Fig.□5F and Table S8). Notably, we observed strikingly coordinated oscillations in the expression of mitochondrially encoded ETC mRNAs (Fig. 5G). These cycling transcripts include the mRNA for ATPase6 (Fig. 5D, Supplementary Fig. 6D) and account for approximately two-thirds of all detected mitochondrial-encoded mRNAs (8 out of 12 in C2; 7 out of 12 in C3, Table S7). In contrast, all cycling mRNAs encoding cytosolic ribosomal proteins peaked near the end of the night in both neuronal subtypes (Fig. 5H). Similarly, mRNAs encoding mitochondrial ribosomal proteins (transcribed in the nucleus) also exhibited a late-night peak in their cycling profiles although with more variability (Supplementary Fig. 6F; see examples in Fig. 5D and Supplementary Fig. 6F). These temporal patterns suggest a coordinated separation between ATP production (during the day) and protein synthesis (during the night), which could be important for preparing the cells for periods of elevated neuronal firing during the light phase.

In addition to ribosomal genes, C2-specific cycling transcripts were enriched for translation-related factors, including *Non1*, *Unr*, *orb2*, *eEF1*α*2*, *eEF2*, *eIF3e*, and *eIF3l* (Figure□5F and Table S8). In C3 cells, the cycling transcript set included genes involved in RNA metabolism (*Cbp20*, *tho2*, *SmB*, *Spf45*) as well as multiple transcriptional regulators and repressors (*sr*, *Hr38*, *fru*, *usp*, *cwo*, *cbt*; Fig.5F and Table S8). *Drosophila* ARGs are not in the GO term databases, but by testing manually, we found that were strongly enriched among cyclers in both cell types (p=7.9e-4 and p=7.78e-4 in C2 and C3 respectively).

### A cell-extrinsic circadian process drives RNA oscillation in C2 and C3 neurons by different mechanisms

To determine if C2 and C3 neurons are under circadian control, we performed the mRNA cyclic analysis in the first day in constant darkness (DD1). We identified 143 and 219 RNA cycling in C2 and C3 cells respectively (Fig. 6A, Table S9), indicating that these neurons receive and integrate circadian information.

**Figure 6:**
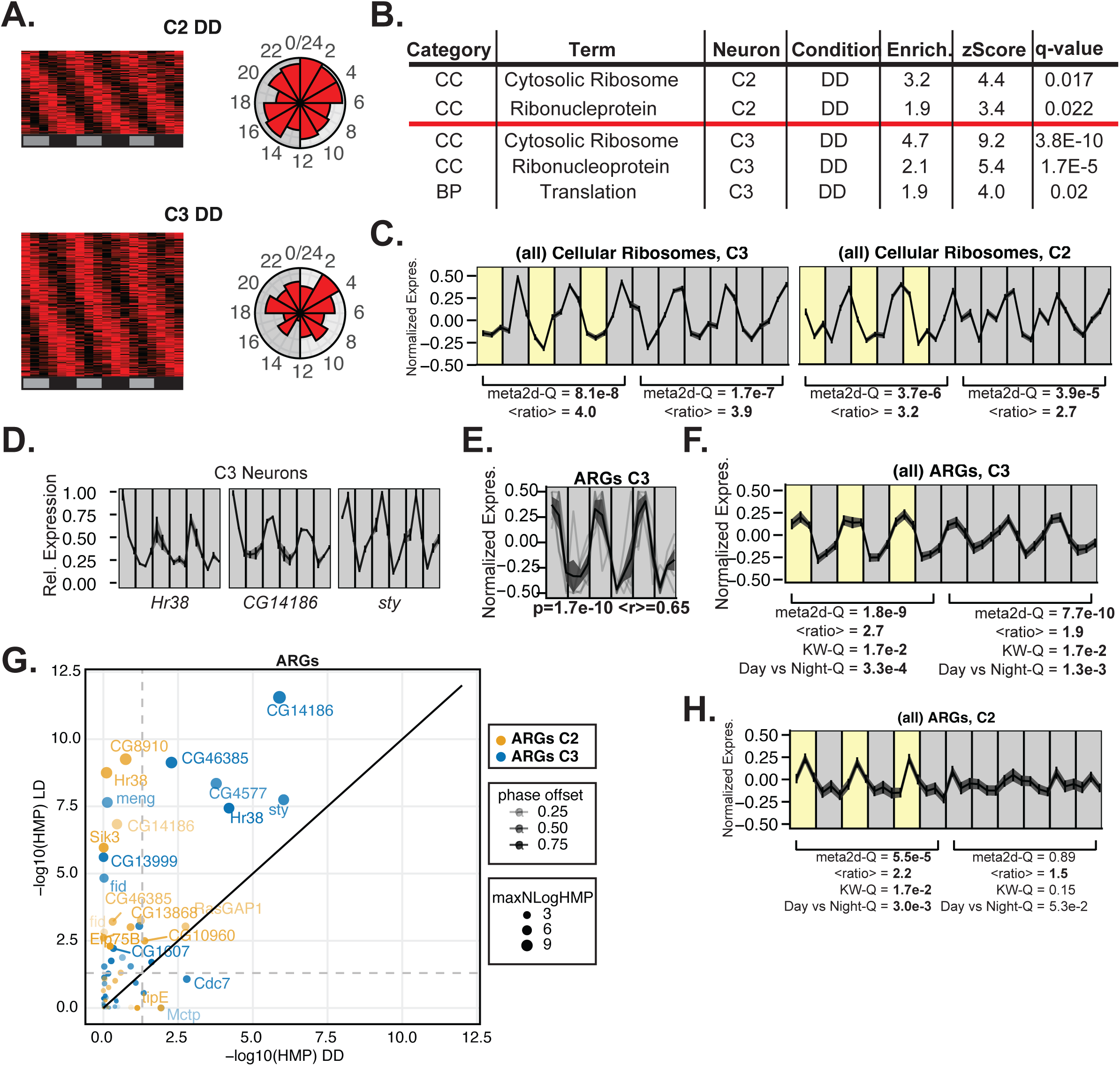
Extrinsic circadian regulation of C2 and C3 neurons drives common and divergent transcriptional responses. **A.** Left: heatmap of scaled gene expression across times in C2 (top) and C3 (bottom) neurons in DD. Right: histograms of inferred peak phases. **B.** Strongest and representative Gene Ontology enrichments for the DD rhythmic mRNAs in C2 and C3 neurons. **C.** Aggregate expression of all expressed mRNAs encoding ribosomes (cellular) along LD and DD. Waveforms are plotted as means ± S.E.M. For each cell type and condition we computed meta2d Q-value of pseudoreplicate means and mean of pseudoreplicate-day expression ratios. **D.** Profiles of cycling mRNAs in C3 in DD, plotted as max-normalized means and S.E.M. among pseudoreplicates. **E.** Aggregate waveforms of cycling ARGs in DD for C3, means ± S.E.M. with individual genes faintly marked. **F.** Aggregate expression of all expressed ARGs along LD and DD in C3 neurons. Waveforms are plotted as means ± S.E.M. For each group we computed: meta2d q-value of pseudoreplicate means, mean of pseudoreplicate-day expression ratios, Kruskal Wallis test q-values among pooled 1-day timepoints, and Wilcoxon test q-values across pooled day vs night. Bold marks q<0.05, mean ratio > 1.5, and mean r > 0.25. **G.** Scatter plot of logged Harmonic means p-values (HMP) among detected activity-regulated genes in DD vs LD. Grey dotted lines mark p=0.05, uncorrected. **H.** Aggregate expression of all expressed ARGs along LD and DD in C2 neurons. Waveforms are plotted as means ± S.E.M and statistics are as described in F.

Both cell types showed significant enrichment for cycling ribosomal protein transcripts (Fig. 6B, Table S10). Remarkably, cytoplasmic ribosomal protein mRNAs displayed widespread coherent oscillations with peak expression during subjective night in both C2 and C3 neurons (Supplementary Fig. 7A and Table S11). This coordination was nearly universal: all (26/26) cycling ribosomal protein transcripts peaked ∼CT22 in C3 cells, and 75% (8/12) in C2 cells (Table S11). The aggregated expression of all ribosomal protein mRNAs (including non-cycling ones) maintained robust 24-hour oscillations with phase and amplitude similar to LD conditions (Fig. 6C). In C2□DD, the aggregate signal of cycling mRNAs encoding cytoplasmic ribosomal proteins exhibits strong cycling and some phase coherence (Supplementary Fig. 7A) yet only a single ARG (RasGAP1) meets cycling criteria (Table S11, Supplementary Fig. 7B). Additional translation-related factors, including initiation factors (eIF3a, eIF3d1, eIF4E1) and release factor eRF1, also cycled in C3 neurons (Fig. 6B).

Analysis of activity-regulated genes (ARGs) revealed distinct mechanisms between the two cell types. In C3 cells, 5 of 37 detected ARGs passed cycling criteria, including *Hr38, CG14186* and *sty* (Fig. 6D, Table S11), a significant enrichment over chance (p=0.004). These cycling ARGs displayed coherent oscillations (Fig. 6E), and the aggregate ARG signal maintained strong midday-peaking rhythms in both LD and DD conditions (Fig. 6F). In DD three out of the four strongest activity-responsive ARGs were expressed (*Hr38*, *CG14186* and *CG46385*). Of these three, two remained significantly rhythmic in C3-DD under FDR correction, with a third (CG46385) showing significant meta2d cycling pvalues but narrowly missing FDR correction (p=0.063, Fig. 6G, Supplementary Fig. 7C, Table S9). These findings suggest that C3 oscillations are driven at least partially by circadian modulation of neuronal activity.

In striking contrast, C2 neurons maintained broad transcriptional cycling in DD despite complete loss of ARG rhythmicity. None of the four strongest ARGs showed significant cycling in C2-DD, and the aggregate ARG signal lost all rhythmic properties, with neither day-by-day nor day-vs-night comparisons reaching significance (Kruskal-Wallis and Wilcoxon tests, Fig. 6H). However, ribosomal protein transcript cycling persisted with amplitude and phase comparable to LD conditions and C3 neurons (Fig. 6C).

These results reveal two distinct mechanisms for extrinsic circadian control: C3 neurons rely on activity-dependent pathways, while C2 neurons use an activity-independent mechanism bypassing activity-regulated pathways. Both mechanisms coordinate ribosomal protein expression, suggesting that circadian regulation of translation machinery represents a fundamental output of the circadian network even in cells without a canonical clock.

### Daily and circadian gene expression is achieved by different mechanisms

Daily and circadian mRNA rhythms *in*□*Drosophila* arise through a spectrum of mechanistically distinct strategies rather than a single, cell-autonomous blueprint. Our integrated single-cell and bulk-tissue analyses reveal four archetypal modes of transcriptional timing that together explain the diverse cycling patterns observed across the nervous system and peripheral organs (Fig.□7).

1. **Autonomous clocks** (Fig. 7A). Some cells, like those of the Malpighian tubule, harbor a complete molecular clockwork (*Clk*/*cyc* activation and *per*/*tim* feedback) and sustain self-sustained oscillations independent of the brain.
2. **Trainable clocks** (Fig. 7B). Other canonical oscillators, such as fat-body cells, retain the core transcriptional loop but rely on the neuropeptide PDF from central pacemaker neurons to entrain their phase, illustrating direct long-range coupling between the brain and peripheral clocks ^99^.
3. **Partial clocks** (Fig. 7C). This group is exemplified by the Lamina wide-field neurons. These cells express only one or few core clock components (e.g. *tim*) yet express high levels of the PDF receptor. These partial clocks might operate with timers rather than real clocks and they could run for one cycle (or half) and damp quickly without reinforcement.
4. **Clockless, driven cells** (Fig. 7D). Certain clock negative cells are timed by alternative cues. C3 neurons, for instance, display hundreds of daily and circadian cyclers, including ARGs, that persist in constant darkness, implicating circadian modulation of neuronal firing. In contrast, C2 neurons maintain rhythmic expression of ribosomal protein and mitochondrial transcripts in DD in the absence of ARG oscillations, indicating an ARG independent, extrinsic mechanism.

**Figure 7:**
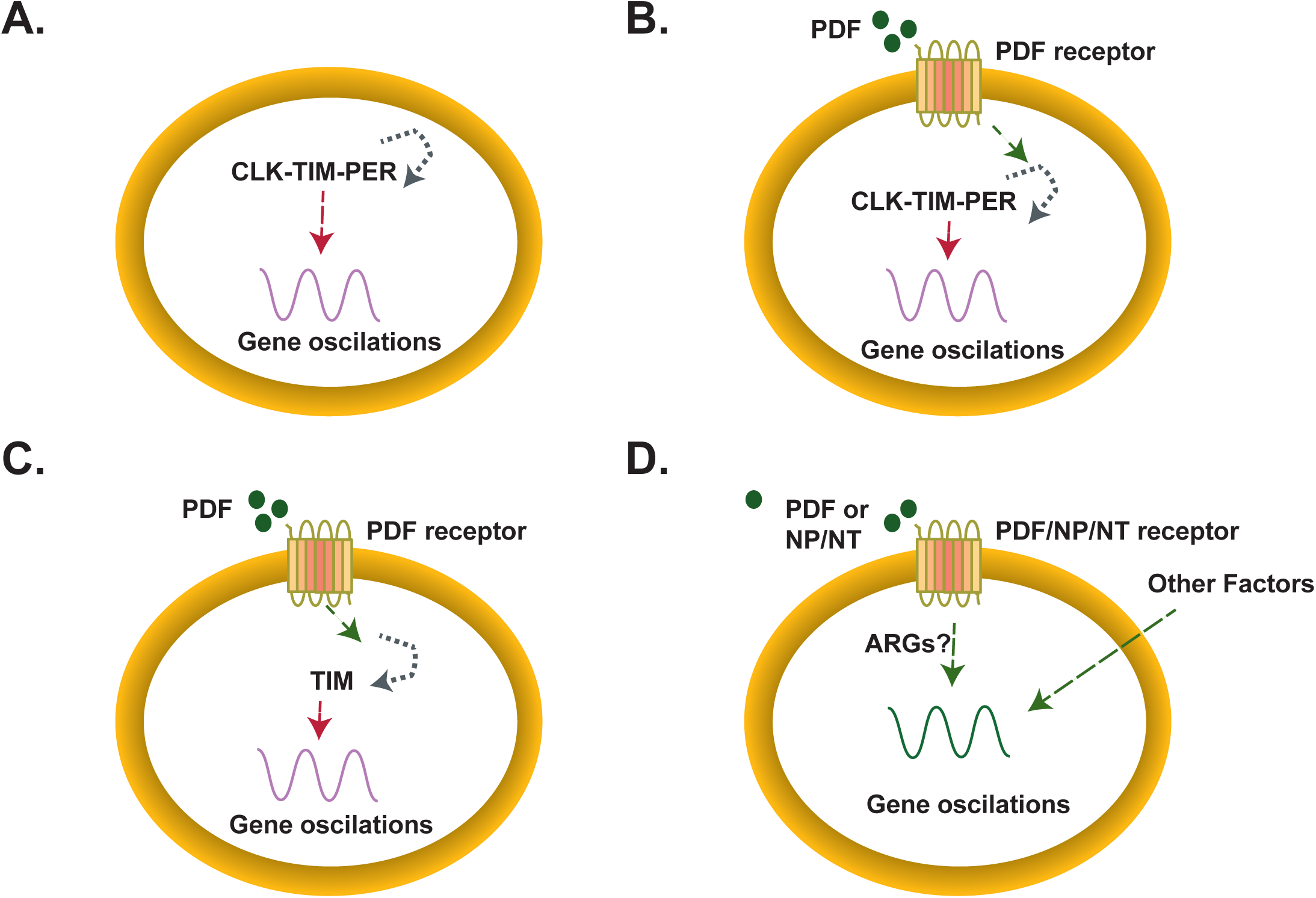
Four different possible models of circadian gene regulation. **A.** Autonomous clocks: some cells, harbor a complete molecular clockwork (*Clk*/*cyc* activation and *per*/*tim* feedback) and sustain self-contained oscillations independent of the brain. These cells have an endogenous clock that is independent of PDF. **B.** Entrainable clocks: other canonical oscillators, such as fat-body cells, retain the core transcriptional loop but rely on the neuropeptide PDF released from central pacemaker neurons to entrain their phase, illustrating direct long-range coupling between the brain and peripheral clocks Cells have an endogenous molecular clock that can be tuned by PDF. **C.** Partial clocks: this group is exemplified by Lamina wide-field neurons in which only one or few core clock genes are expressed. In addition, they express high levels of the PDF receptor. **D.** Clockless, driven cells: these cells do not have an internal molecular clock but express high levels of *Pdfr* or another neuropeptide or neurotransmitter receptor, suggesting that the central pacemaker could regulate the observed oscillations in gene expression through paracrine signaling or direct synaptic contact.

Several general principles emerge. First, high amplitude mRNA cycling can occur in the absence of a complete feedback loop, challenging the binary “clock positive versus clock negative” view. Second, remote regulation by neuropeptides or synaptic input is widespread; nearly every cycling, clock negative cluster expresses at least one circadianly relevant neuropeptide receptor. Third, distinct extrinsic signals sculpt different transcriptome modules: PDF imposes a dawn biased program, activity linked cues drive midday ARG peaks in C3, and an unidentified mechanism coordinates late night ribosomal and mitochondrial mRNA surges both in C2 and C3. Collectively, these observations demonstrate that the circadian clock distributes temporal information through multiple, partially overlapping channels, thereby permitting rhythmic physiology and behavior across a far broader cellular landscape than previously appreciated.

## DISCUSSION

By systematically mapping core clock gene expression and interrogating daily transcriptional rhythms at cell-type resolution, we uncovered a surprisingly intricate landscape of circadian regulation in *Drosophila*. While earlier work focused on canonical pacemaker neurons, glia and photoreceptors, our data reveal that rhythmic gene expression pervades many additional cell types, even those that lack a complete set of core clock genes. This expanded repertoire of circadian regulation goes beyond the traditional view of cell-autonomous oscillators and suggests temporal control as an emergent property of a distributed neuronal and peripheral network. Our findings illustrate that circadian rhythms are governed not by a single, uniform mechanism, but by a hierarchy of oscillators and signals. In addition to cell-autonomous clocks, many cells achieve rhythmicity through extrinsic timing cues, effectively, a multi-tiered network architecture reminiscent of the mammalian SCN’s core-loop and output hierarchy.

Our findings complement previous reports of intrinsic clocks in peripheral tissues, such as the fat body and Malpighian tubules ^100,101^, and parallel observations in mammals, where lung epithelia, macrophages, and other peripheral cells harbor circadian oscillators ^102–105^. We extend this concept by demonstrating that diverse *Drosophila* cell types, including tracheal, epithelial, and haemocyte-like cells, exhibit core clock gene expression, highlighting a widespread distribution of circadian timing across tissues. This broad landscape of oscillators is consistent with the idea that temporal regulation is integrated throughout the organism.

Our analysis also exposes discrepancies with physiological studies that reported circadian responses in olfactory, gustatory and other chemosensory neurons ^67,106–108^. We failed to detect *Clk*, *per* or *tim* enrichment in any of these sensory cells. There are several non□exclusive explanations for this discrepancy. First, the single□cell datasets we employed were harvested at undefined zeitgeber times, so we may have missed transient peaks of low□expressed transcripts. Second, the rhythmic sensory outputs could arise from clock genes translated from undetected mRNA pools. Finally, these neurons might rely entirely on extrinsic cues from *bona fide* clock cells rather than on intrinsic oscillators. Time□resolved single□cell profiling will be indispensable for addressing these possibilities.

Among the most provocative findings is the existence of neurons that express only subsets of the canonical transcription-translation circadian loop, thereby forming “partial” oscillators. Lawf1 and Poxn neurons illustrate this intermediate state: Lawf1 neurons display high amplitude *tim* cycles in both LD and DD with undetectable *Clk* or *per*, whereas Poxn expresses *per* in the absence of *tim* or *Clk*. Such partial clocks challenge the dogma that a complete feedback loop is an absolute requirement for circadian timing. This raises the possibility that cells can operate as ‘timers’: oscillators that perhaps run for one cycle or damp quickly without reinforcement, rather than self-sustained clocks. Such timers might provide local rhythmic cues yet remain dependent on the central pacemaker for long-term synchrony. Functional dissection by targeted knockdown of *tim* in Lawf1 will clarify whether these neurons influence behavioral outputs or align to central pacemakers through PDF or other neuromodulators. Notably, emerging connectomics studies ^39^ indicate that central clock neurons extend broad projections via both synaptic and neuropeptidergic pathways, consistent with our observation of extrinsic entrainment in different cell types.

Our findings converge with accumulating evidence that circadian timing is governed by a hierarchy of cues. To our knowledge, this is the first direct evidence of circadian output (persisting in constant darkness) in neurons that lack all known core clock genes. While there is a recent report of sorted Mushroom Body neurons from *Drosophila* brains showing RNA rhythms in DD ^44^, the bulk sorting approach makes it impossible to determine whether the observed cycling occurs in the neurons themselves or in contaminating clock-containing cells (e.g., glia). Indeed, we observed expression of some clock genes (*per* and *Clk*) in these data. Our single-cell approach confirms that truly clockless neurons can still be driven to oscillate by network signals. The presence of *PdfR* in many clockless cell types suggests that PDF released by central pacemakers can entrain downstream targets observe that approximately half of the ∼230 robust brain cyclers peak around ZT□21, mirroring the known phase of PDF release, at least in LD ^32^. Consistent with our finding previous work demonstrated the influence of PDF to diverse neuronal circuits. For example, PDF signaling onto non-clock dopaminergic PPM3 neurons inhibits them during the day promoting daytime wakefulness ^41^. PDF also acts on DN1p clock neurons to induce DH31 release at dawn, which ultimately drives morning arousal ^109^. These examples illustrate how a neuropeptide can orchestrate neural circuits beyond the canonical clock network. This PDF-driven regime is particularly evident in the visual system, where neurons in the accessory medulla region ^45,46^, provide light information to LNvs and LNds while linking back to photoreceptors. Similarly, cells such as L2 neurons, which show rhythmic changes despite low *per* levels ^53,110^, highlight that timing signals can propagate through unconventional pathways to shape daily physiological states. The extensive wiring reported by Reinhard *et*□*al.* ^39^ provides a plausible scaffold for such multi-synaptic dissemination of timing cues.

Our single-cell circadian experiments on C2 and C3 neurons expose two distinct modes of extrinsic control. Both neuron types lack the core clock molecular machinery yet exhibit hundreds of cycling transcripts. In C3, ARGs such as *Hr38* and *CG14186* remain rhythmic in DD, and their phases align with midday peaks in LNv calcium ^88^, suggesting that circadian modulation of neuronal activity drives these oscillations. Conversely, C2 neurons lose ARG rhythms in DD but retain coherent cycles of ribosomal and mitochondrial mRNAs, implying an alternative, ARG-independent mechanism. The presence of the *Pdfr* suggests that these cells receive signals from clock cells which could then modulate second-messenger cascades driving gene expression rhythms. The prominence of ribosomal protein (RP) and mitochondrial electron transport chain (ETC) transcripts among cyclers in both C2 and C3 raises intriguing metabolic considerations. Daytime peaks of ETC mRNAs coincide with anticipated high ATP demand, whereas nocturnal surges of RP mRNAs may prime the translational apparatus for early-day activity. Similar temporal partitioning of bioenergetic and biosynthetic processes has been reported in mammalian liver and heart ^111–114^, suggesting conserved principles.

Our study has limitations. The single-cell atlases we mined lack temporal metadata, obliging us to infer clock status from averaged expression profiles and mitigating phase bias by gene scaling. Although our bulk RNA-seq confirmed many single-cell predictions, comprehensive time-series single-cell sequencing will be required to capture low-amplitude or phase-restricted transcripts and validate clock status conclusively. Moreover, we examined only PDFR among neuropeptide receptors; expanding the repertoire to include AstA-R1, sNPFR, NPF-R and others ^42,100,115,116^ will refine our map of extrinsic circadian communication.

Our results support a model in which circadian timing in *Drosophila* is orchestrated by a mosaic of autonomous oscillators, partial clocks and remotely driven cells. Central pacemaker neurons, exemplified by the LNvs, act not merely as master clocks but as hubs that transmits temporal information through synaptic, peptidergic and activity-dependent channels. Peripheral tissues and diverse neuronal classes interpret these cues via cell-specific receptor repertoires, second-messenger pathways and transcriptional circuits, tailoring rhythmic programs to local physiological demands. Such distributed architecture echoes the mammalian hierarchy in which the SCN entrains subordinate tissue clocks via neural and humoral signals, yet peripheral oscillators retain flexibility to respond to feeding, temperature and other systemic cues ^117–120^.

Recognizing that circadian outputs can emerge from full molecular loops, partial feedback modules or purely extrinsic inputs reframes how we think about clock disruption in disease and ageing. It also raises the possibility that therapeutic manipulations of timing signals could restore rhythms in cells where the circadian clock has deteriorated with age. Future work should establish how diverse extrinsic cues converge on specific transcriptome modules and determine if similar hybrid mechanisms operate in mammalian brain circuits and peripheral organs.

## MATERIALS AND METHODS

### Fly stocks and husbandry

*w*^1118^ and UAS-eGFP (referred to as UAS-GFP) fly strains were obtained from the Bloomington *Drosophila* Stock Center (Indiana, USA). The split GAL driver targeting the C2 and C3 neurons ^121^ was obtained from the Janelia FlyLight collection (SS00779: R20C11-p65ADZp in attP40; R48D11-ZpGdbd in attP2). Panel of strains with a nuclear GFP reporter (UAS-H2A-GFP) and carrying 3^rd^ chromosomes from wild-type strains (DGRP, *Drosophila* Reference Genetic Panel ^122^ were previously described in ^123^, Lawf1 and Lawf2-Gal4 lines (1118-Gal4 and 11D03-Gal4) were previously described ^124^. The *tim-*Tomato reporter line was generated in our lab and previously described ^125^. All crosses were performed and raised at 25 °C in a 12:12 Light/Dark cycle (LD) conditions.

### Immunohistochemistry

Lawf2-Gal4 lines were recombined with the UAS-GFP line to generate the *w*^1118^; UAS-GFP; Lawf2-Gal4 line. *tim*-Tomato reporter flies consist of a codon-optimized and destabilized tdTomato fluorescent protein, which has three copies of a nuclear localization signal (NLS) to focus signals for quantification under the control of the *tim* promoter (12). *w*^1118^; UAS-GFP; Lawf2-Gal4 line was crossed to the *tim*-Tomato reporter line. +; UAS-GFP/*tim*-Tomato; Lawf2-Gal4) males were selected from their progeny. Three to five days old flies were anesthetized with CO_2_, their whole body fixed in 4% PFA for 1 hour at room temperature (RT), and their brains were dissected in PBS at ZT0, ZT4, ZT8, ZT12, ZT16, and ZT20. Brains were subjected to a standard immunostaining protocol. Briefly, the brains were incubated overnight at 4 °C in a mix of mouse anti-GFP (1:1000, Sigma) and rabbit anti-DsRed (1:1000, Rockland) and 7% normal goat serum in PBS. Samples were washed in PBS and incubated with anti-mouse Alexa 488 (1:500, Jackson ImmunoResearch) and anti-rabbit Cy3 (1:500, Jackson ImmunoResearch) for 2 hours at RT, washed in PBS and mounted in anti-fade mounting medium Fluoromount-G (Southern Biotech).

Single snapshots for each brain were obtained using a Zeiss LSM 880 confocal microscope. Image analysis was performed in Fiji (ImageJ, NIH-USA). Statistical analysis and data visualization was performed in Prism 8 (GraphPad).

### In Situ

All *in situ* procedures were performed using RNAscope Multiplex Fluorescent Detection Kit v2 (Advanced Cell Diagnostics, ACD) as described in ^126,127^ with few modifications. Briefly, *Drosophila* brains were dissected at ZT3, ZT15, CT3 and CT15 directly into 4% formaldehyde in PBS and fixed for 16 hours at RT. Following fixation, brains were pretreated as described in ^127^. Brains were incubated with prewarmed Dm-tim-C1 probe (Catalog # 1323121-C) overnight at 40° C. Signal amplification steps were performed according to manufacturer’s instructions followed by incubation in Vivid 650 dye (Tocris bioscience) at 1:1500 dilution in TSA buffer for 30 minutes at 40 o C. Following RNAscope, brains were stained with rabbit anti-GFP (Abcam, ab290), as done in ^127^. Z-stacks covering the whole optic lobe were obtained using a Leica Stellaris 8 confocal microscope. Image analysis was performed in Fiji by generating masks that tracked individual cells to measure the intensity of *tim* signal within each (ImageJ, NIH-USA). Statistical analysis and data visualization were performed in Prism 8 (GraphPad).

### Generation of bulk RNAseq datasets

*w*^1118^ flies were entrained for at least 3 days at 25 °C and dissected at six different timepoints (ZT3, ZT7, ZT11, ZT15, ZT19, and ZT23). RNA from the fly brains was extracted using TRIzol reagent (Sigma, T9424) and treated with DnaseI (NEB, M0303L). We used 150 ng of RNA as input for preparing 3’ RNA sequencing libraries following CelSeq2 protocol (100,101), changing the UMI to 6 bases. Sequencing was performed on Illumina NextSeq 500 system. The data was deposited in GEO (GSE233184).

### Single Cell Sequencing of C2/C3 neurons

The C2/C3 split-Gal4 line was crossed to a panel of strains expressing nuclear GFP (UAS-H2A-GFP) and carrying different wild-type third chromosomes with known genotypes ^123^. Samples from each timepoint and replicate were tagged using a chromosome from a unique DGRP strain ^122^. Flies were entrained in LD (12:12) for 3 days, with DD flies then subjected to 1 day of constant darkness. Brains were dissected at target timepoints and dissociated into a single-cell suspension using enzymatic dissociation ^52^. All timepoints and replicates from were pooled and processed together as a single sample. GFP-expressing cells were purified using BD Melody fluorescence-activated cell sorter, and cell concentration was estimated using a hemocytometer. The single-cell suspension was used to generate one scRNA-seq library using the Chromium Single Cell 3□v3 kit. The resulting library was sequenced using the Illumina HiSeq4000 system (2 x 150 bp reads) by Novogene. The data was deposited in GEO (GSE297050).

### Computational Analysis

#### Single-cell RNA processing of publicly available data

Already published single cell pre-processed data was downloaded from the respective GEO databases. For the fly cell atlas ^128^, an in-house script was written to integrate the data into Seurat format (https://github.com/ipatop/FlyCellAtlas.download.with.annotations). Briefly, we gathered the data and performed the normalization and clustering necessary to access the complete list of normalized genes. Cell type assignment was extracted from the published data. We validated the cell-type assignment by manually inspecting marker genes. The fly cell atlas datasets ^128^ come from dissected tissues except for the fat body, coenocyte, and tracheal cells that were FACS sorted using a tissue-specific GAL4 driving UAS–nuclear-GFP. The brain single cell dataset we utilized was GSE116969. For the optic lobe dataset ^57^, we downloaded the data from GEO (GEO: GSE156455). Integration and clustering steps were needed to process this dataset. We used the parameters reported by the authors.

For enrichment and clustering analysis, gene expression was retrieved using the *FetchData* function from Seurat. Gene level sum was summarized by cell type using general R functions. After that, z-scaling and centering were done first over the gene level and then over the cell type level. Heatmaps and clustering were done using the ComplexHeatmap package. The optimal k-mean cluster number was determined using the *eclust* function with 500 randomization steps. All functions used for this analysis are stored in the following R-Package: https://github.com/ipatop/SingleCell_SumScaledExpresion.git

#### Sorted cell-type RNA processing

Data was downloaded from GEO (GEO: GSE103772 ^74^) and aligned to the *Drosophila melanogaster* dm6 genome and transcriptome version using STAR ^129^. Quantification was done using with feature counts. Mean normalized counts were used with ComplexHeatmap for clustering and visualization. The optimal k-mean cluster number was determined using *eclust* function with 500 randomization steps.

#### Circadian gene assessment from bulk RNA sequencing

For the 3′ RNA-seq, data were aligned to the *Drosophila melanogaster* dm6 genome and transcriptome version using STAR ^129^, and quantification was done with ESAT ^130^. The circadian analysis was performed using the package MetaCycle ^95^. For each circadian timepoint, three replicates were analyzed. Since each replicate was sequenced separately, the counts were divided by the maximum value in each replicate after normalizing by library size. Genes with more than two zero counts at any time point were discarded. The amplitude for each replicate was calculated as the maximum divided by the minimum for each gene. We used the MetaCycle ^95^ to apply the JTK algorithm which we used for circadian analysis (105). A gene was considered a cycler if the JTK p-adjusted value was less than 0.05 and the amplitude was more than 1.5.

#### Analysis of C2/C3 single cell data

##### Data processing and analysis

Sequencing reads were processed using the standard Cell Ranger pipeline from 10X Genomics (version 7.1.0). The reference genome and gene annotations were obtained from FlyBase (release 6.29, ^131^). Single cells from each sample were demultiplexed using demuxlet (version 2, https://github.com/statgen/popscle, ^94^) as described in ^123^. Demultiplexing was performed using genotypes of 24 experimental strains and an extra 7 strains as negative controls. Out of 4805 cell barcodes, 4376 (91%) were assigned to the correct genotypes, and others were classified as doublets and ambiguous barcodes (none were assigned to the 7 control genotypes). The analysis was conducted using the standard Seurat 4.1.3 workflow ^132^. The high-quality transcriptomes were selected using the following criteria: assigned to the expected genotypes; unique genes/cell: 200–4,000, mitochondrial transcripts < 10%. Unsupervised clustering was performed using 12 principal components, a clustering resolution of 0.2, and otherwise default settings. Two large clusters were annotated as C2 (1943 cells) and C3 (2089 cells) neurons based on known marker genes ^57^; 131 cells were removed from further analysis. The final dataset was evenly distributed and had a high coverage across all samples, with a minimum of 115 cells for each cell type and timepoint. To filter non-expressed genes, for each timecourse (LD or DD) we extracted the percentage of cells expressing a given gene using the DGRP repliactes as a second set of 6 timepoints. Genes with >10% in at least two timepoints were considered expressed in this cell type and condition. This threshold was chosen after assessing expected gene contamination as a function of expression level of literature markers for photoreceptors and glia: *ninaE*, *Rh2*-*Rh6*, *trp*, *trpl*, *ninaA*, and *HisCl1*; *repo*, *alrm*, *wrapper*, *Hml*, *moody*, *Indy*, *Vmat*, *e*, *pnt*, *ttk*. The final dataset with cell type annotations, experimental conditions, and individual DGRP genotypes are available at NCBI GEO (GSE297050). Analysis was performed via a custom script (https://github.com/rashkovert/C2C3/).

##### Analysis of cycling transcripts

Similar to ^48^, we subsampled data per time point (per cell-type and condition) into three pseudoreplicate days, and assessed 24-hour (periods of 20hr to 28hr) rhythmicity using the MetaCycle v1.2.0 R package ^95^. To reduce variation in rhythmicity detection among possible subsamplings, we performed this process three times and for each gene we combined the p-values using the method of harmonic means ^48^. To further eliminate noise we applied a second filter (max/min values >1.5) and then corrected the pvalues using the Benjamini-Hochberg procedure. We defined genes as cycling when the FDR<0.05. To obtain the final phase of oscillation we averaged the values obtained in the 3 meta2d outputs in polar space. ***GO terms analysis.*** To identify enriched biological processes among circadian cycling genes, we performed Gene Ontology (GO) enrichment analysis using the cluster Profiler R package (v4.8.1, ^133^) in conjunction with the org.Dm.eg.db annotation database for *Drosophila melanogaster* ^134^. Enrichment analysis was performed using the enrichGO function, specifying keyType = “FLYBASE” (FBgn identifiers), and evaluating each of the three GO ontologies: Molecular Function (MF), Biological Process (BP), and Cellular Component (CC). Parameters included minGSSize = 5, maxGSSize = 500, and multiple testing correction using the Benjamini–Hochberg (BH) method. Cycling gene sets for each neuronal subtype and condition (C2_LD, C3_LD, C2_DD, C3_DD) were tested against matched background gene sets corresponding to all expressed genes in each context. Only GO terms with at least 5 overlapping genes were retained, and false discovery rate (FDR) values were recalculated using p.adjust() to obtain q.recalc. GO terms with q.recalc < 0.10 were considered significant.

##### Cycling assessment of groups of cycling genes

Original gene groupings were taken from gene ontology annotation using org.Dm.eg.db (v3.19.1). Activity-regulated genes were taken from ^98^, using genes found to be induced by at least 2 of 3 experiments. We defined the “strongest ARGs” as those that exhibited >2 fold-change upregulation within at least 2 experiments. To determine enrichment of ARGs among cycling genes we generated using Fisher’s exact test relative to the set of expressed genes per mode/condition. To determine if a given group of cycling genes display coherent cycling, we scaled their expression around the mean, average them and scaled to the maximum. This averaged expression was used to determine a cycling p-value using the meta2d function of the package MetaCycle. For determining phase coherence among cyclers within a group we performed Pearson correlation between the expression of this gene across the timecourse and the average waveform of the group of genes. We then averaged the correlation values across the group and obtained a final correlation index named <r> in the graph.

##### Assessment of collective cycling

To determine whether the expression of a gene group displays collective oscillations, we averaged their normalized and scaled expression across the timecourse to generate an averaged/scaled waveform. We then performed meta2d analysis to determine cycling, in addition to determine differences between day and night we used the Mann-Whitney U-test and as an additional measurement of difference among timepoints we used the Kruskal-Wallis test.

### Statistical Methods

The utilized test and statistical methods are detailed across the analysis in the method section, and the utilized tests are also reported in the Fig.Legends.

## Data and materials availability

The datasets generated and/or analyzed during the current study are available in the GEO repository. The data generated in this study has been deposited in GEO (accessing number GSE233184 for the bulk RNAseq data and GEO GSE297050 for the single cell data).

## Acknowledgments

We thank Dr. Claude Desplan for the Lawf2-Gal4 fly lines, Aljoscha Nern for the C2/C3 split Gal4 line and Dr. Sagiv Shifman for his insight on the data analysis processing.

## Author contributions

Conceptualization: SK, ILP, AMA, TR

Investigation: AMA, ILB, NB, KC

Data Analysis: ILP, TR, NB, ILB, YK

Supervision: SK, MFC, YK

Writing—original draft: SK, ILP, AMA

Writing—review & editing: SK, ILP, AMA, TR, MFC, NB, YK

## SUPPLEMENTARY INFORMATION

**Supplementary Fig. 1.**
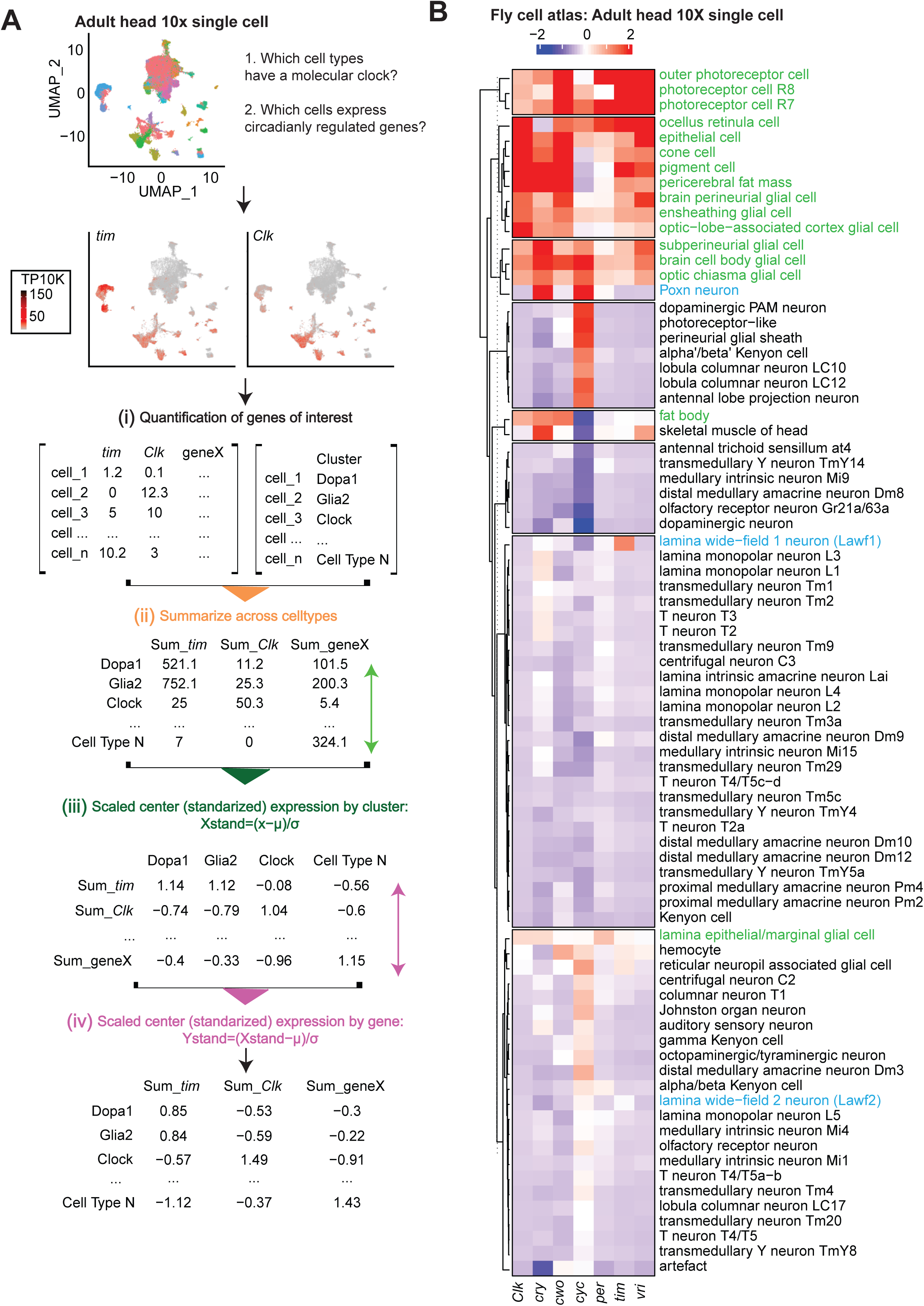
Standardized gene expression recapitulates known core-clock expression pattern in the head. **A.** Approach to identify whether a cell type expresses clock genes. Top: Representative UMAP plot from the fly head single cell dataset colored by cell type assignment and showing *tim* and *Clk* gene expression. Red indicates higher expression (color bar, TP10K). Bottom: Representation of the processing pipeline used for analysis: i) Generate the matrix of raw counts of gene expression *per* cell type; ii) Calculate the sum expression of each gene by cell type; iii) Standardize the expression values across clusters; iv) Standardize those expression values across genes. We performed this analysis for the following 10X single cell datasets: Brain *(72)*, Optic Lobe *(56)*, and Fly cell atlas *(128)*. **B**. Heatmap of standardized values of core-clock genes from Head 10X data with blue and red representing low and high levels, respectively. Clusters are calculated with k means (data: *(128)*).

**Supplementary Fig. 2.**
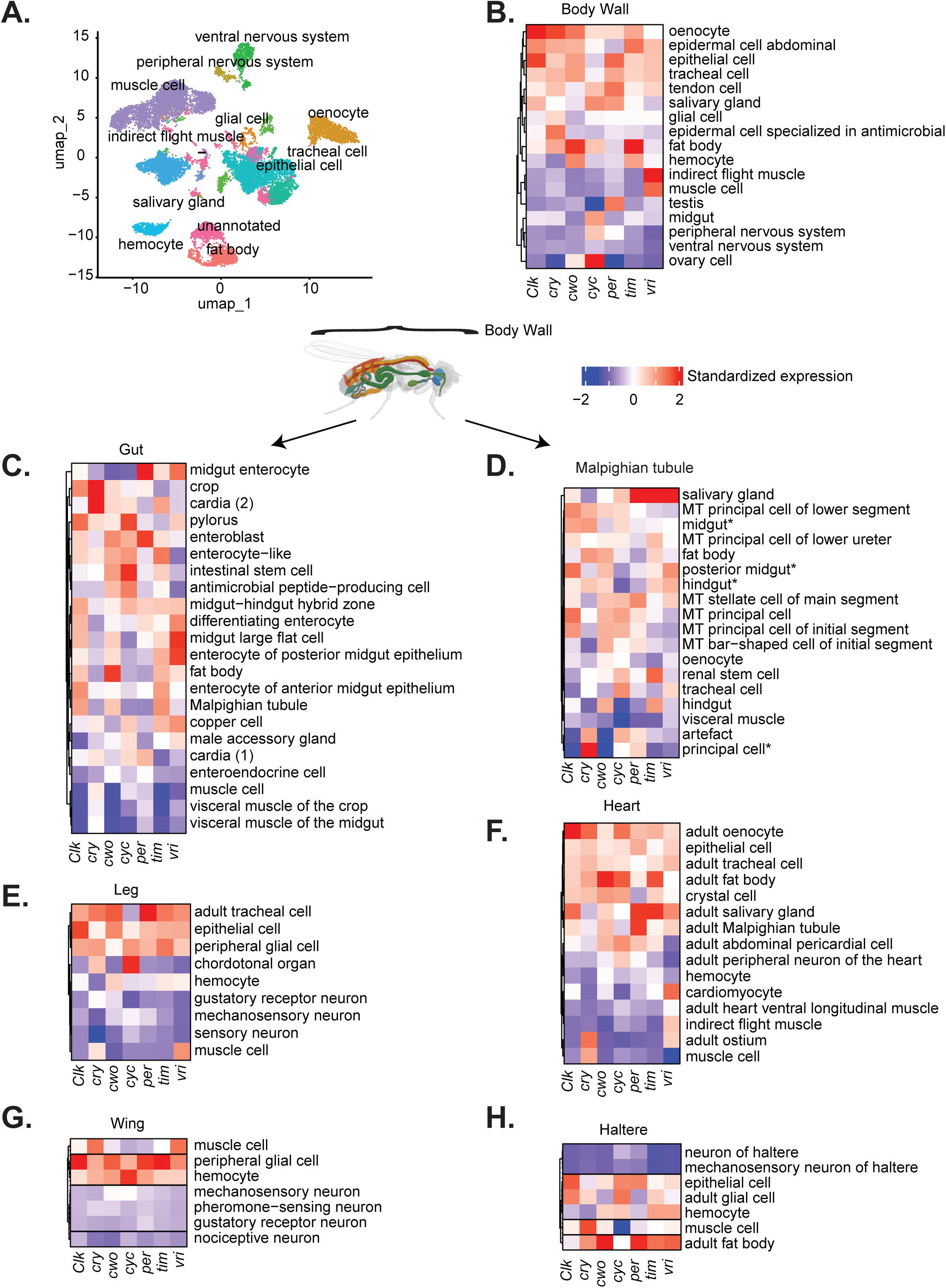
Cell types across the body have diverse core-clock enrichment patterns. **A.** Representative UMAP of body wall 10X single-cell data colored by cell type assignment. **B-H**. Heatmaps of standardized values of core-clock genes from 10X data of the body wall, gut, Malpighian tubule, leg, heart, wing, and haltere in *Drosophila*. Blue and red represent low and high expression levels, respectively. Clusters are calculated with k means (data: *(128)*).

**Supplementary Fig. 3.**
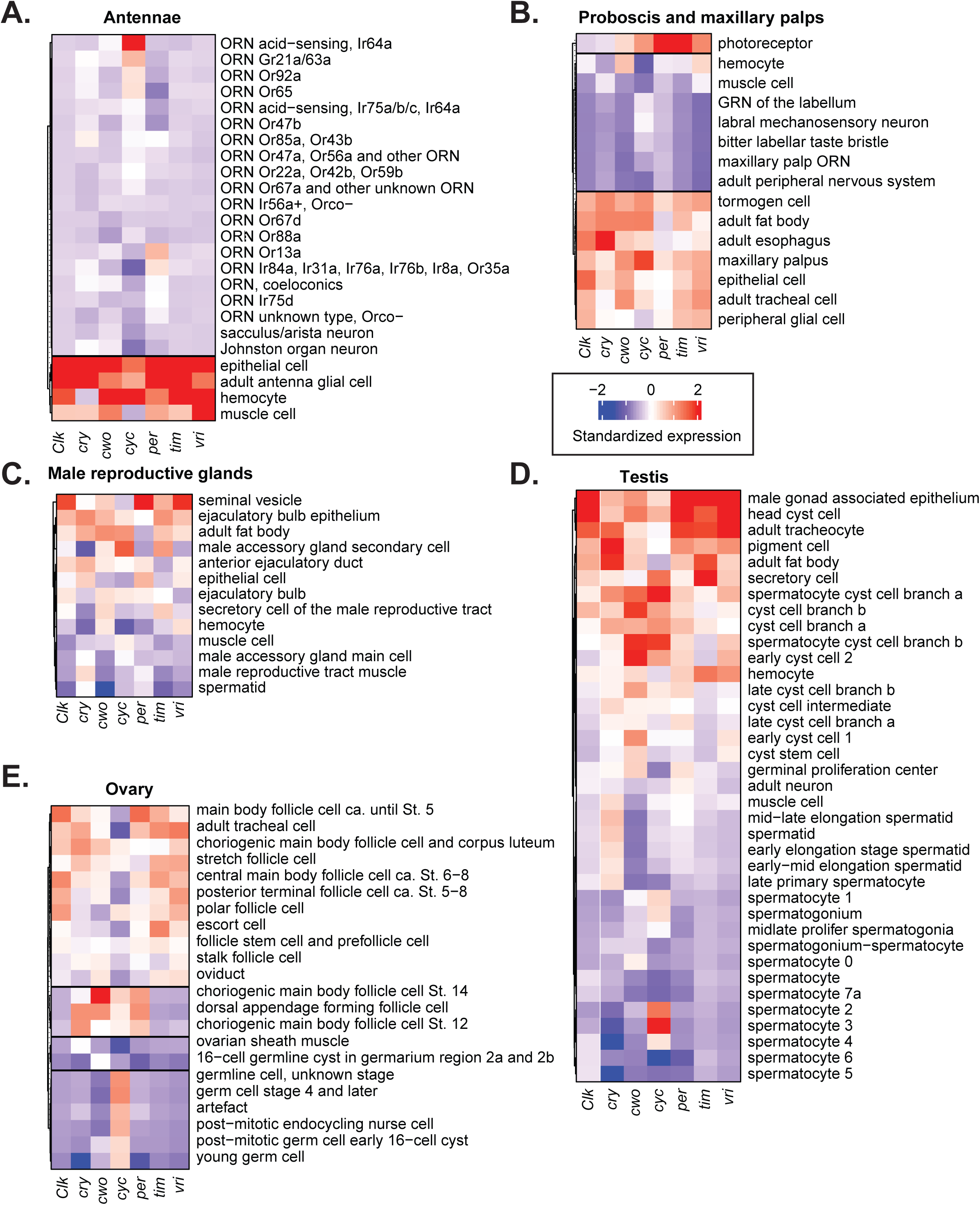
Cell types across the body have diverse core-clock enrichment patterns. **A-E:** Heatmaps of standardized values of core-clock genes in Antennae, Proboscis and maxillary palps, Male reproductive glands, Testis and Ovary in *Drosophila*. Blue and red represent low and high expression levels, respectively. Clusters are calculated with k means (data: *(128)*).

**Supplementary Fig. 4.**
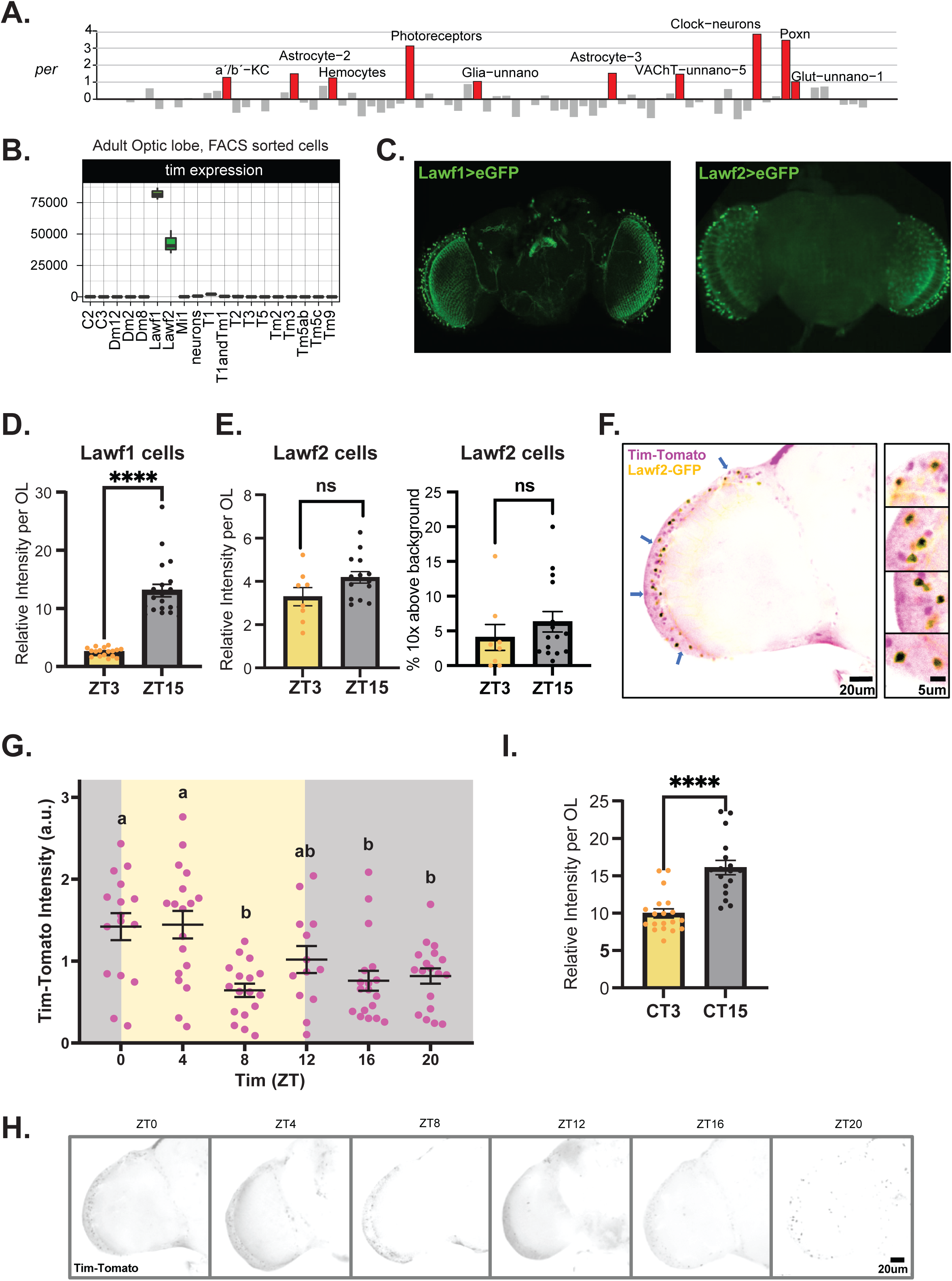
*Tim* is highly expressed and display daily changes in Lawf neurons. **A.** Standardized *per* expression from 10X single-cell data from the brain. In red, cell types with expression levels higher than one standard deviation from the mean (data: *(72)*). **B.** Boxplot showing normalized *tim* expression in FACS sorted cells from the optic lobe (data: *(73)*). **C.** Expression of the Lawf1 and Lawf2 Gal4 driver and overlap with *tim* expressing cells. Top: Lawf1 (left) or Lawf2 neurons (right). Cells were visualized by GFP in Lawf1(1)-Gal4 (or Lawf2-Gal4); UAS-eGFP flies. Center: *tim* expression as visualized by *in situ* hybridization using probes against *tim* coding region. Image was taken at ZT15 timepoint. Bottom: overlay of the two images. **D.** Quantification of *tim* mRNA fluorescence intensity in Lawf1 neurons at ZT3 and ZT15 per optic lobe. For each optic lobe, Intensity was calculated as the mean *tim* mRNA fluorescence in all Lawf 1 neurons. Mann Whitney test, p value <0.0001. **E.** Quantification of *tim* mRNA fluorescence intensity in individual Lawf2 neurons at two timepoints (ZT3 and ZT15). Left: Quantification per optic lobe. For each optic lobe, intensity was calculated as the mean *tim* mRNA fluorescence in all Lawf 1 neurons. Right: Proportion of Lawf 1 neurons per optic lobe with *tim* mRNA signal ≥10-fold above background. Mann Whitney test was not significant (ns) in both cases. **F.** *w*^1118^; UAS-GFP/*tim*-Tomato; Lawf2-Gal4 fly brain immunostained with anti-GFP (yellow) and anti-dsRed (magenta) antibodies. Blue arrows indicate the cells that are shown in magnified photos on the right. The regions where both signals overlap are marked in black. **G.** *tim*-Tomato intensity quantification in Lawf2-GFP expressing cells across time points. ZT (zeitgeber time) being ZT0 and ZT12 the time at when lights turn ON and OFF, respectively. Each dot on the plot represents the mean intensity value for a single brain. ZT0 = 1.422 + 0.165 (n=16), ZT4 = 1.445 + 0.169 (n=18), ZT8 = 0.643 (n=17) + 0.082, ZT12 = 1.019 + 0.165 (n=13), ZT16 = 0.760 + 0.122 (n=18), ZT20 = 0.818 + 0.094 (n=18); N=2. One way ANOVA followed by Bonferroni’s multiple comparisons test, F(5,94) = 6.691, p< 0.0001. Only different letters (but not the combination of them) indicate significant differences: p<0.05 for ZT0 vs ZT16, ZT0 vs ZT20, and ZT4 vs ZT20; p<0.01 for ZT4 vs ZT16, and ZT0 vs ZT8; p<0.001 for ZT4 vs ZT8. Data expressed in Mean+SEM. **H.** Representative images of *tim*-Tomato brains immunostained with anti-dsRed antibody dissected at the indicated time points. **I.** Quantification of *tim* mRNA fluorescence intensity in Lawf2 neurons at two circadian timepoints (CT3 and CT15). For each optic lobe, Intensity was calculated as the mean *tim* mRNA fluorescence in all Lawf 1 neurons. Mann Whitney test, p value <0.0001.

**Supplementary Fig. 5.**
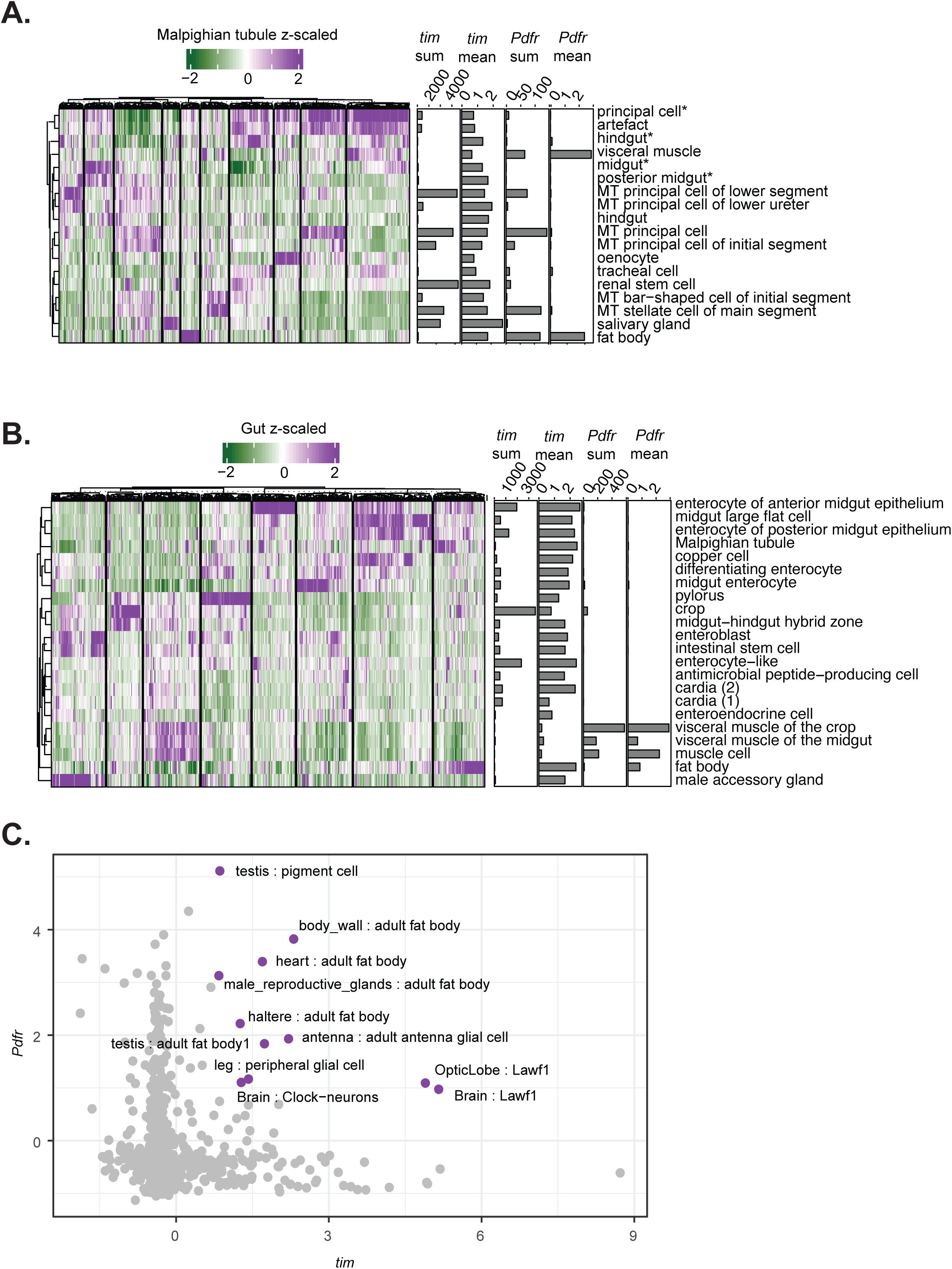
Expression of cycling genes in peripheral tissues correlate with high levels of core clock genes or *Pdfr*. **A.** Heatmap of enrichment of cyclic genes in the Malpighian tubule in single-cell clusters. In bar plot, sum and mean expression of *tim* and *Pdfr* for each cluster. **B.** Heatmap displaying enrichment of genes that cycle in the gut in single-cell clusters. **C.** Standardized expression of *tim vs. Pdfr.* In violet, the clusters with high *tim* and *Pdfr* scaled expression.

**Supplementary Fig. 6.**
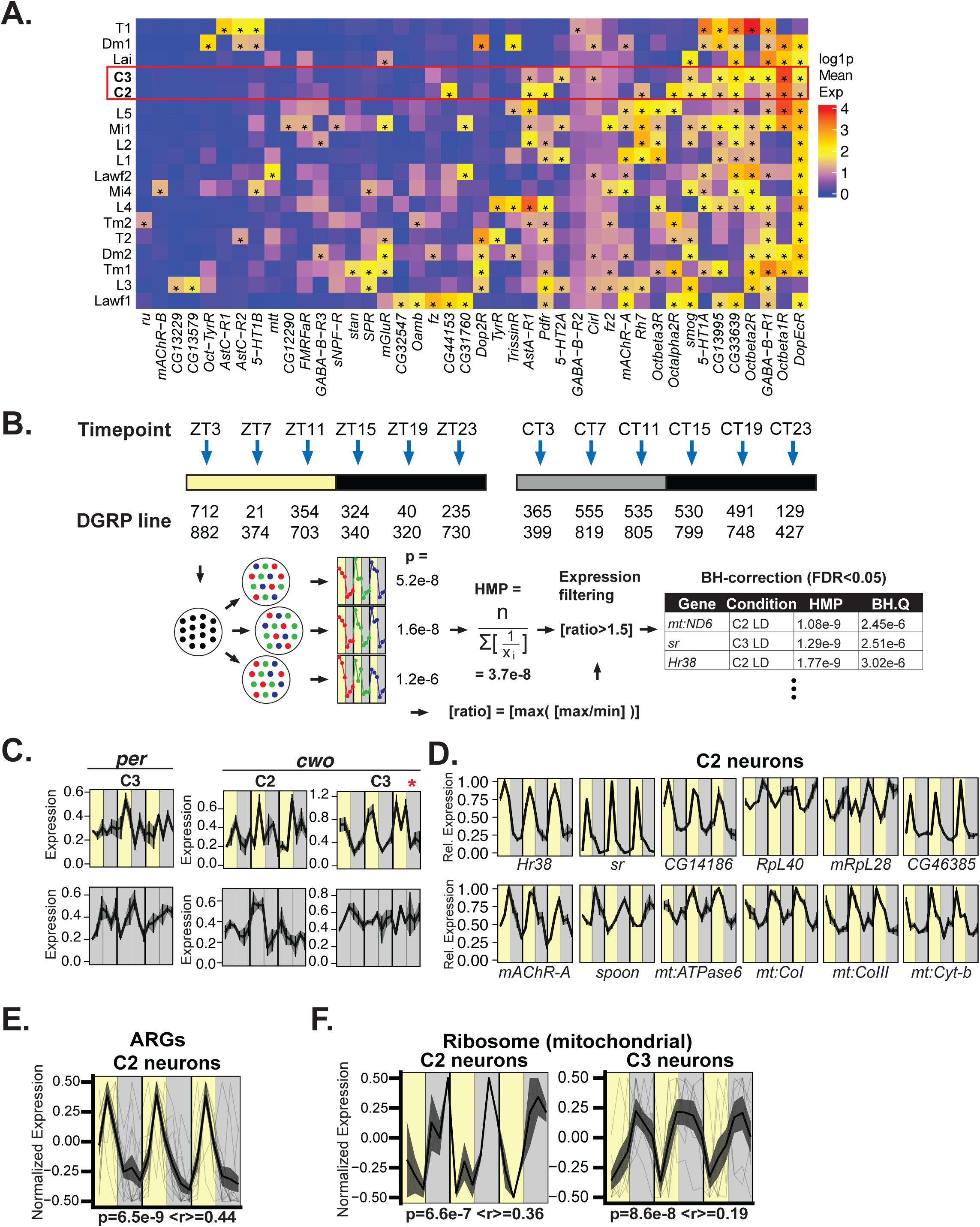
C2 and C3 neurons exhibit rhythmic and coordinated transcript expression. **A.** Heatmap of GPCR gene expression (GPCRs from *(91)*) of highly expressed (mean expression > 2 – marked by asterisks) transcripts in select optic cell types (data from *(72)*). **B.** Schematic of circadian sc-RNAseq experiment and analysis. Flies of duplicate DGRP lines were entrained to LD or DD at 6 collection times. Cells were randomly assigned to one of three pseudoreplicate-days three times, and meta2d p-values were combined under assumed non-independence by the harmonic means method *(95)*). Ratio-wise filtering was applied before Benjamini-Hochberg correction across genes to FDR<0.05. **C.** Expression of canonical clock genes *per* and *cwo* in LD and DD. Waveforms are means+S.E.M. of pseudoreplicates. * denotes that *cwo* was identified as cycling in LD. **D.** Cycling gene profiles of representative mRNAs in C2-LD, plotted as max-normalized means with S.E.M. bars+ribbons among pseudoreplicates. Results include ARGs, mitochondrial transcripts (ETC), and ribosomal mRNAs. **E.** Aggregate waveforms of cycling ARGs in C2 neurons. The waveform was calculated by averaging the scale expression of the genes within the group (see methods). Faint lines mark individual genes, dark lines indicate signal averages, ribbons mark S.E.M. The waveforms were rescaled to -0.5 to 0.5 by subtracting 0.5. p indicates meta2d p-value and <r> the average of pairwise Pearson’s correlations between the individual genes and the average waveform. **F.** Similar to E but for mRNAs encoding mitochondrial ribosomal proteins in C2 (left) and C3 (right) neurons.

**Supplementary Fig. 7.**
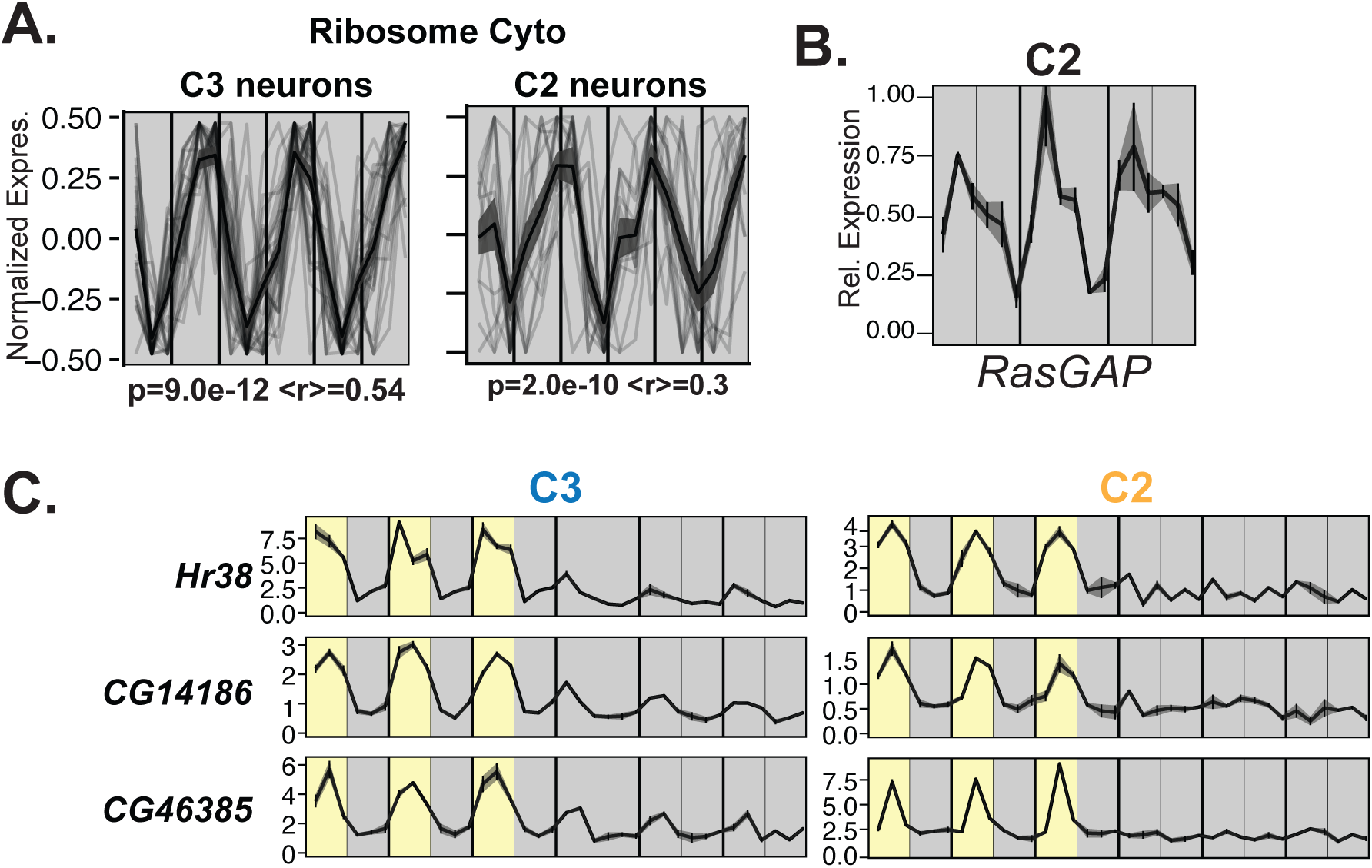
Extrinsic circadian regulation of C2 and C3 neurons regulates common and divergent transcriptional responses. **A.** Aggregate waveforms of cycling ribosomal protein transcripts (cellular), in DD for C2 neurons. The waveform was calculated by averaging the scale expression of the genes within the group (see methods). Faint lines mark individual genes, dark lines indicate signal averages, ribbons mark S.E.M. The waveforms were rescaled to -0.5 to 0.5 by subtracting 0.5. p indicates meta2d p-value and <r> the average of pairwise Pearson’s correlations between the individual genes and the average waveform. **B.** Cycling gene profile of *RasGAP* in C2 neurons in DD, plotted as max-normalized means and S.E.M. bars±ribbons among pseudoreplicates. **C.** Expression of the indicated ARGs along LD and DD in C2 (right) and C3 (left) neurons. Waveforms are plotted as means ± S.E.M.

## Supplementary Table Legends

**Table S1. Standardized levels of *Clk*, *Pdfr*, *cry*, *cwo*, *cyc*, *per*, *tim*, and *vri* in all single-cell datasets (data from Davie et al 2018, Konstantinides et al 2018, Kurmangaliyev et al 2020, and Li et al 2022)**

**Table S2. Sum expression of each core clock gene by cell type and percentage of contribution to the total summed signal in the fly brain (data from Davie et al 2018).**

**Table S3. Normalized gene expression in different cell types of the optic lobe sorted by Fluorescence-activated cell sorting (data from Konstantinides et al 2018)**

**Table S4. Cycling parameters of genes with daily oscillations in a 6-timepoint experiment under 12 hours of light-dark cycles (FDR from JTK algorithm < 0.05 and minimum amplitude > 2, data from this study).**

**Table S5. Enrichment of cycling genes in the different cell types identified in single cell data**

**Table S6. Expression and Cycling analysis of RNAs in C2 and C3 cells in LD**

**Table S7. Definition of gene groups and their detection and cycling status in LD in C2 and C3 cells**

**Table S8. Gene Ontology (GO) terms enriched among cyclers in C2 and C3 neurons in LD.**

**Table S9. Expression and Cycling analysis of RNAs in C2 and C3 cells in DD**

**Table S10. Gene Ontology (GO) terms enriched among cyclers in C2 and C3 neurons in DD.**

**Table S11. Definition of gene groups and their detection and cycling status in DD in C2 and C3 cells**

